# A national baseline for methane sink habitats and methanotroph diversity

**DOI:** 10.64898/2026.02.02.703227

**Authors:** KS Knudsen, M Sereika, TBNJ Jensen, F Delogu, T Schmider, C Jiang, RH Kirkegaard, AT Tveit, PH Nielsen, M Albertsen, CM Singleton

**Author notes:** Corresponding AuthorCaitlin M. Singleton, Center for Microbial Communities, Department of Chemistry and Bioscience, Aalborg University, Fredrik Bajers Vej 7H, 9220 Aalborg, Denmark.

## Abstract

Methane emissions account for nearly a third of the Earth’s effective radiative forcing, with methanotrophs playing a critical role in mitigating emissions by oxidising methane in diverse environments^1^. Despite their ecological importance, methanotrophic diversity and environmental distribution remain incompletely characterised due to cultivation challenges, incomplete or low-quality metagenome-assembled genomes, and limited taxonomic resolution in marker gene surveys. Here, we present a national study of the biogeography of novel and known methanotrophs across Denmark’s major natural, urban and agricultural habitats, using genome-resolved classification of 10,683 metagenomes^2^ and 102 new methanotrophic species^3^. By linking metabolic potential to habitat-specific distributions, we reveal uncharacterised methanotrophs as dominant in natural ecosystems. These findings provide a comprehensive baseline of methanotroph diversity, reveal clear contrasts between natural and disturbed habitats, and highlight candidate species and habitats for future methane-mitigation strategies.

## Main

Methane is a potent greenhouse gas, with increased emissions accounting for approximately 31% of the effective radiative forcing on Earth since pre-industrial times^1^. Around 70% of methane emissions originate from methanogenic archaea in natural and anthropogenic systems such as wetlands, peatlands, landfills, and agriculture^4,5^. Much of this methane (50-80%) is consumed before reaching the atmosphere by methanotrophs, including aerobic methane oxidising bacteria (MOB) and anaerobic methanotrophic archaea^6^. Additionally, specialised methanotrophs (atmospheric methane oxidising bacteria (atmMOB)) are able to oxidise methane at its atmospheric concentration (∼1.93 ppm), representing the only known biological sink of atmospheric methane. Approximately 10% of the global sink for atmospheric methane is ascribed to atmMOB activity in upland habitats, with similar contributions from tree surfaces and soils (∼30 Tg year ^-1^, each)^1,7^, but this sink is impaired by human activities such as deforestation and intensified land use^8^. Characterisation and isolation of methanotrophs from high-methane environments has provided insight into oxidation kinetics, substrate specificity, and metabolic pathways. However, the cultivation process is selective, isolation efforts often rely on elevated methane concentrations, and methanotrophs contributing to the atmospheric methane sink have proven difficult to obtain in pure culture^9–11^.

To assess environmental methanotroph diversity, studies have often relied on the amplification of the marker genes *pmoA* or *mmoX*, encoding subunit A of the particulate (pMMO), or subunit X of the soluble (sMMO) forms of the methane monooxygenase, respectively^12,13^. The recurrent detection of *pmoA* sequence clusters distinct from characterised methanotrophic lineages in upland soils with net methane consumption highlights the lack of ecologically relevant atmMOB isolates^8,14^. Investigations into uncultured groups are complicated by the differences in both taxonomic resolution and capture bias between primers^12,15,16^, making comparisons across studies difficult. Additionally, methane monooxygenases share homology with ammonia and hydrocarbon monooxygenases^17–19^ increasing the risk of functional misinterpretation. Recent advances in the recovery of metagenome-assembled genomes (MAGs) from complex environments, such as soil^3^ allows us to circumvent these issues by combining the interpretation of marker genes within the broader genomic context, including genes for the initial steps of methane oxidation, downstream energy conservation and carbon assimilation.

Here, we link functional marker genes, genomes and genomic potential to provide a detailed taxonomic classification of environmentally relevant aerobic methanotrophs across Denmark using the Microflora Danica dataset. This Atlas of Danish microbes comprises 10,683 short-read metagenomes^2^ and 154 deep long-read metagenomes^3^, from which we describe the diversity and biogeography of canonical and novel MOB using gene-and genome-resolved approaches. We reveal habitat specific patterns and provide insight into natural methanotroph communities with methane sink potential. By determining MOB diversity, novelty, and community differences in natural and managed ecosystems, we provide knowledge to guide national and global efforts to improve land management and restoration practices.

### Database curation and targeted sequencing of methanotrophs

As the initial step for the country-wide classification and analysis of Danish methanotrophs, we updated existing *pmoA* and *mmoX* databases with sequences from the increasing availability of high quality (HQ) genomes in the genome taxonomy database (GTDB r220)^20^. We screened GTDB r220 using GraftM^21^ *pmoA* and *mmoX* search packages^22^, and linked *pmoA* and *mmoX* marker genes with taxonomic identity and inferred metabolic potential (**Supplementary Note 1**). We evaluated methane associated metabolism based on presence of modules for methanol oxidation (*mxaF*-type methanol dehydrogenases (MDHs) or *xoxF*-type MDHs), and modules for oxidation of formaldehyde to formate (tetrahydromethanopterin pathway (H_4_MPT) or thiol-dependent (glutathione (GSH)-linked) pathway). Additionally, we assessed the presence of carbon-assimilation modules for formate incorporation into the serine cycle (tetrahydrofolate pathway (H_4_F)), formaldehyde incorporation into the ribulose monophosphate pathway (RuMP) and for CO_2_ fixation via the Calvin-Benson-Bassham pathway (CBB)^4,23^.

We identified canonical methanotrophs encoding pMMO and/or sMMO in GTDB r220 (**Table 1**), but also discovered additional putative methanotrophs across several uncharacterised families with unconfirmed methane oxidation capabilities, for which we examined metabolic potential, gene synteny and MAG contamination (**Supplementary Note 1**). Potential methanotrophs encoding putative pMMOs and further modules of the methane metabolism and carbon assimilation included members of the families DRLZ01 and UBA1147 within the *Methylococcales* order, and the alphaproteobacterial *Skermanella aerolata* (**Table 1, Supplementary Note 1, Extended Data Figure 1, Extended Data Figure 2**). Potential sMMO containing methanotrophs were identified within the families *Xanthobacteraceae* (genus U87765) and *Acetobacteraceae* (genera *Rhodopila* and *Acidiphilum),* and the order Burkholderiales (genera *Methylotenera,* JABFRO01 and AVCC01) (**Table 1, Supplementary Note 1, Extended Data Figure 1**).

**Table 1:**
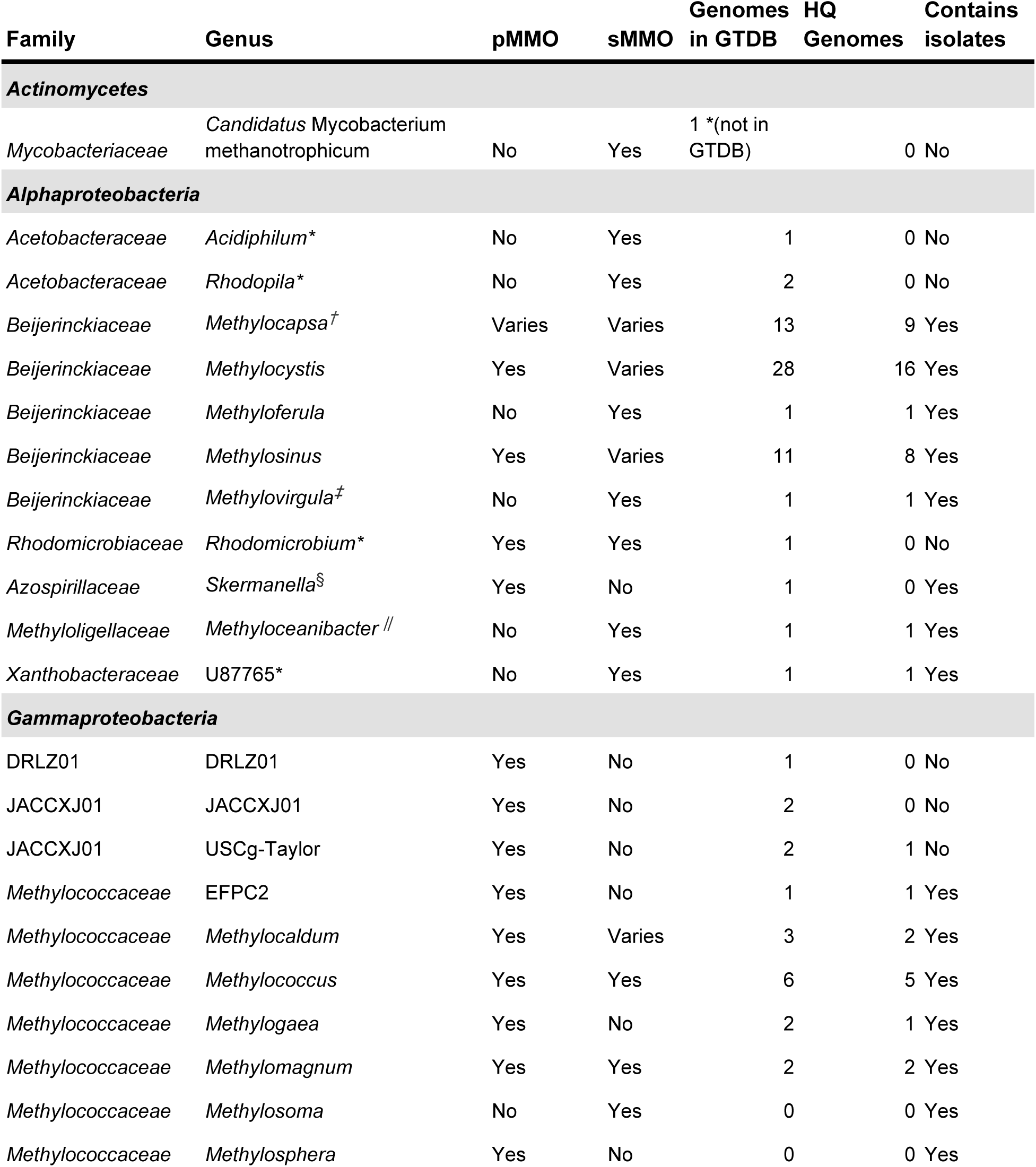
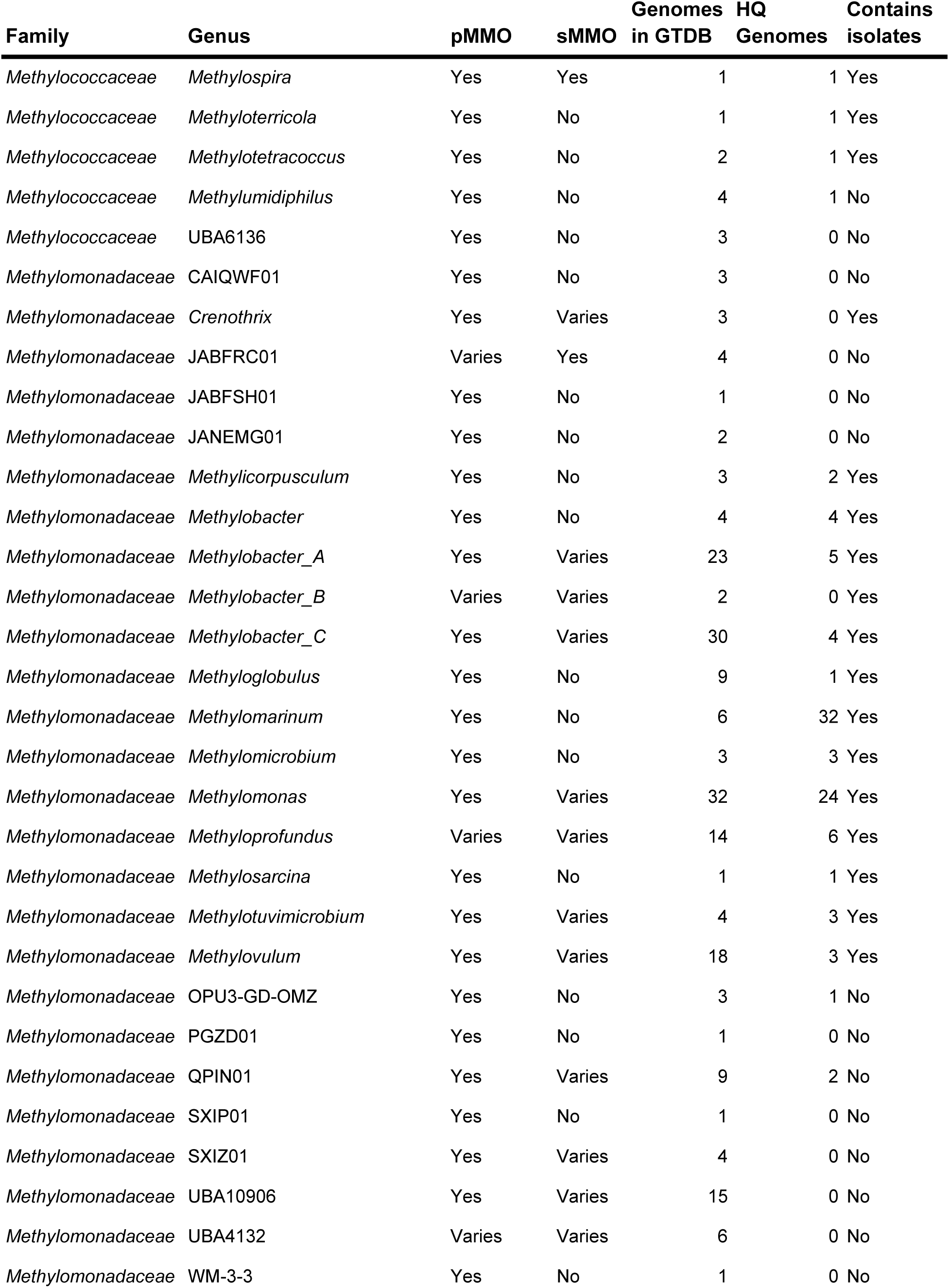
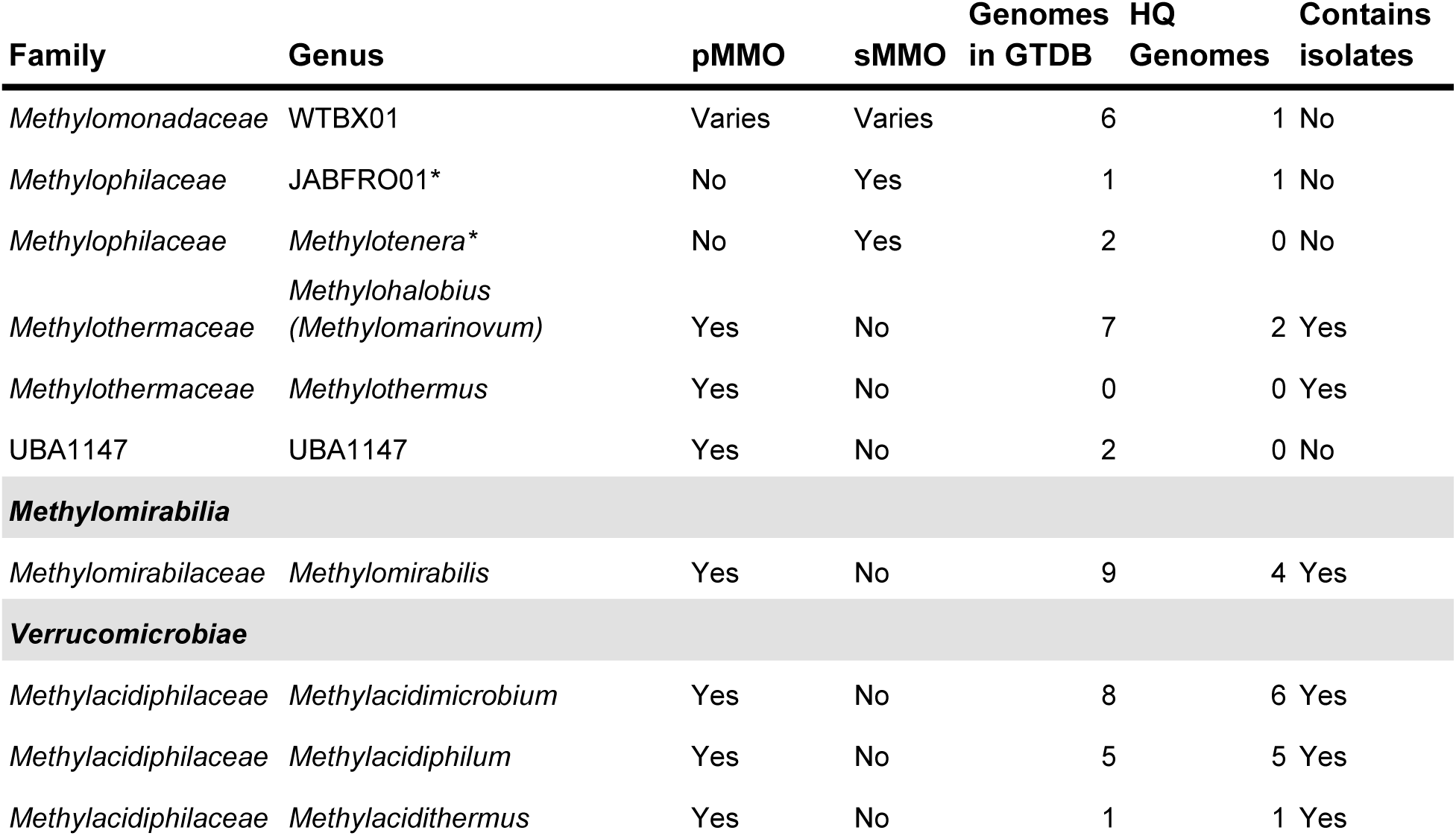
Overview of methane oxidising bacteria and the corresponding type of methane monooxygenase based on GTDB r220 and available literature^12,34–46^. Genomes in GTDB are reported by the number of genomes within each genus that contains at least one type of MMO. Class is highlighted in bold and grey. pMMO: particulate methane monooxygenase. sMMO: soluble methane monooxygenase. HQ: high quality MAGs encoding MMO, according to MIMAG standards (Bowers et al. 2017). *\**Only a subset of MAGs within the genus are putative methanotrophs. *^†^Methylocella* has been renamed to *Methylocapsa* between r220 and r226 of GTDB. *^‡^*Only *Methylovirgula thiovorans* is a methanotroph, while other species within *Methylovirgula* are methylotrophs. ^§^Only *Skermanella aerolata* is a putative methanotroph. *^||^*Only *Methyloceanibacter methanicus* is a methanotroph, while other species within *Methyloceanibacter* are methylotrophs.

Next, we linked putative *pmoA* sequences associated with atmospheric methane oxidation to genomes. *pmoA* sequences associated with atmospheric methane oxidation include those identified in upland soils, such as the frequently observed upland soil clusters (USC), USCα and USCy, and the less frequently occurring *Methylocystis*-like *pmoAs*^13,14,24^. Members of *Methylocystis* have been shown to maintain methane oxidation activity for long periods (*Methylocystis* spp. DWT, LR1^25^, SC2 (*M. hydrogenophila*^26^)). More recently, filter-cultures of *Methylocystis rosea* SV97 and *Methylocapsa* spp. (*M. aurea* KYG, *M. gorgona* MG08 and *M. palsarum* NE2) were shown to grow on air and maintain oxidation of atmospheric methane after 12 month incubations with air as the only carbon and energy source^27^. Only two isolates (*M.* gorgona and *Methylocapsa* D3K7), encode the USCα-like *pmoAs* which are frequently detected in forest soils with atmospheric methane consumption^8,14,28^. In addition to these two species, six additional medium quality (MQ, MIMAG standards^29^) *Methylocapsa* species representatives encoding USCα *pmoA* are present in GTDB r220 (**Table 1**).

For USCγ and tropical USC (TUSC), atmospheric methane oxidation is suspected, but experimental confirmation and isolation of a representative strain is missing^12,24,30^. Within the gammaproteobacterial family JACCXJ01 (proposed *Candidatus* Methyloligotrophales^31^), we identified USCγ *pmoA* sequences in the two genera USCg-Taylor (proposed *Candidatus* Methyloligotropha) (genome accessions: GCA_002007425.1, GCA_030859565.1) and JACCXJ01 (GCA_013697045.1, GCA_030860485.1) (**Table 1, Extended Data Figure 2**). While all genomes encoded methanol and formaldehyde oxidation genes, the key genes distinct to formaldehyde assimilation using the RuMP cycle (*hps, psi*), were missing (**Supplementary Note 1, Extended Data Figure 2**). Recent work links TUSC sequences to the phyla Gemmatimonadota (*Ca.* M. kingii)^32^ and Binatota (Desulfubacterota_B in GTDB r220, previously part of Deltaproteobacteria)^33^. However, in the *Ca.* M. kingii MAG, the *pmoCAB* is located on a small contig (7 kbp, cov: 7x) with high sequence similarity to Binatota (91%, 94%, 95% amino acid identity against nr, 2025-10-28, blastx), while the CDS on other contigs (median cov: 11.4x, mean cov: 11.8x) were assigned to Gemmatimonadota (**Extended Data Figure 3b**). In addition to the potential misbinning of TUSC *pmoCAB* in the MAG, the serine cycle was incomplete, missing the key genes *sga* (serine glyoxylate aminotransferase) and *hprA* (hydroxypyruvate reductase) (**Extended Data Figure 3a**), and alternative methane derived carbon-assimilation pathways could not be identified. Thus, the proposed “*Ca.* M. kingii” is likely not a methanotroph. The gene phylogeny of the putative *pmoA* sequences in Binatota MAGs appeared to be divided into a *bmoA* (butane monooxygenase) clade, and a TUSC-affiliated clade, with some members displaying genomic potential for downstream methane processing^33^ (**Supplementary Note 1**). Whether TUSC sequences encode methane specific monooxygenases remains uncertain.

While several putative methanotrophic lineages were recovered in GTDB r220, some marker gene clades remain poorly connected to genomes, either lacking plausible genome assignments (e.g. TUSC), missing HQ representatives (e.g. USCγ, *Rhodomicrobium*) or represented by few genomes (e.g. USCα). To resolve these issues, we searched 10,683 MFD metagenomes^2^ to identify promising samples for methanotroph MAG recovery, which were then selected for deep sequencing^3^. This search identified 26 promising samples for HQ MAG recovery (**Figure 2b**), targeting the poorly represented USC methanotrophs, the putative *Rhodomicrobium* methanotrophs and the unresolved Binatales and TUSC members (**Figure 1**, **Figure 2a**). Additionally, we aimed to recover MAGs encoding divergent copies of the pMMO, where the in situ role and function remains unclear, namely pXMO (*pxmA*)^47^ and a paralogue of pMMO (*pmoA2*) found in some members of *Methylocystis* and *Methylosinus*^26^ (**Figure 1**, **Figure 2a**). As a result of the targeted sequencing, we recovered 286 medium-to high quality MAGs that encoded at least one copy of *mmoX* or *pmoA*. These represented 115 species (95% average nucleotide identity (ANI)), 102 of which were new. Among them were 34 *Methylocapsa*, 20 *Methylocystis* and seven *Rhodomicrobium* spp. (from 41 new *Rhodomicrobium* spp. recovered) (**Figure 1**). All MAGs encoding USCα-like *pmoA* belonged to *Methylocapsa.* We recovered MAGs encoding USCγ-like *pmoA* within USCg-Taylor (*Ca.* Methyloligotropha) (two new species) and JACCXJ01 (one new species), and TUSC-like *pmoA* encoded MAGs within the Binatales (seven new species). Moreover, we recovered novel species within *Methylomirabilis* (two new species), Methylobacter_A (four new species) and Methylobacter_C (six new species) (**Figure 1**). Overall, we added 82 new full length *pmoA* and 21 *mmoX* sequences to our search databases, tailoring our analysis to the major habitat types of Denmark.

**Figure 1:**
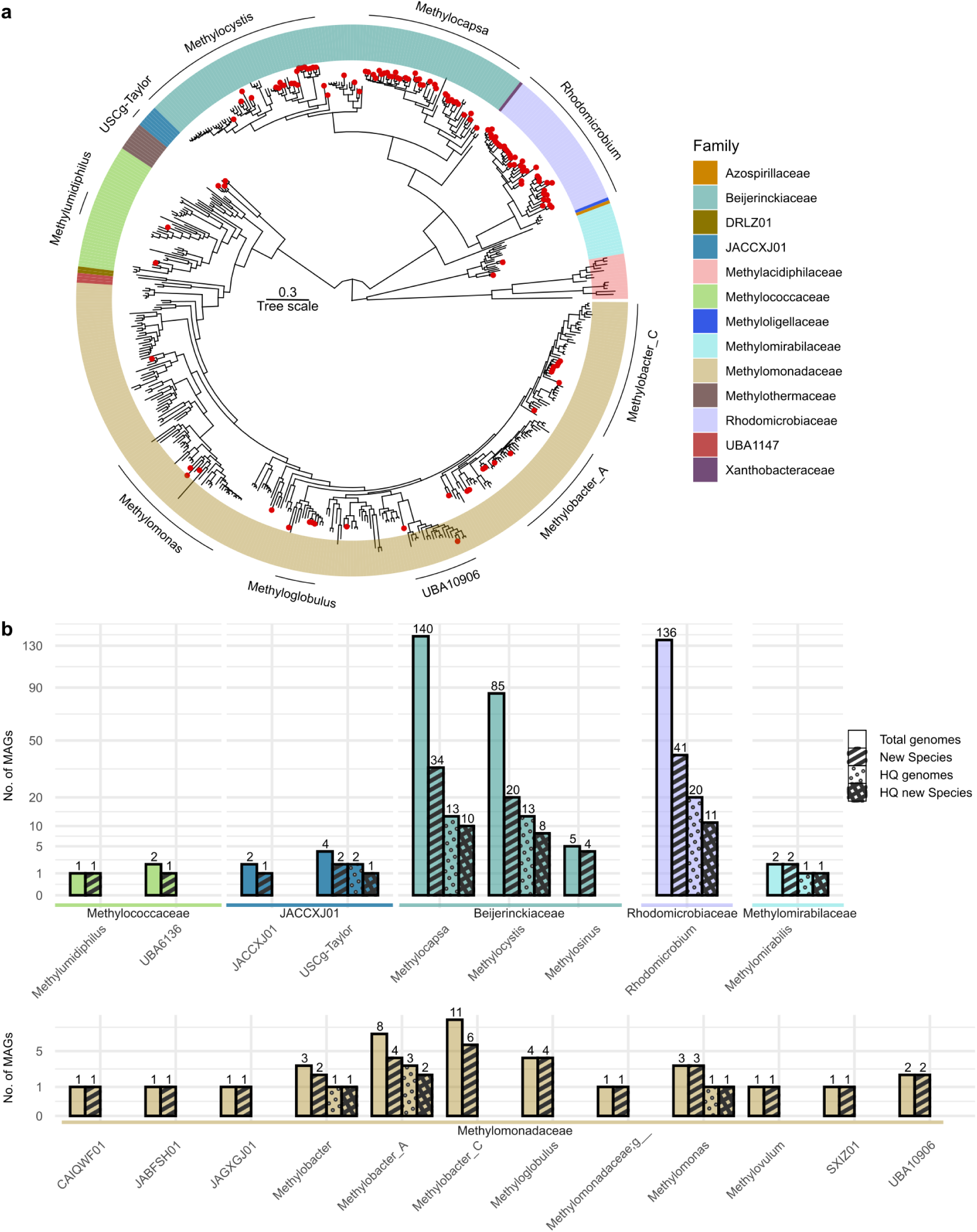
Genome recovery of methanotroph MAGs. **a.** Phylogenomic tree of methanotrophs. Genomes recovered for novel species within genera of putative methanotrophs are indicated with red dots. The bar surrounding the tree indicates the gtdb-tk classification of the clade. Genera of particular interest are indicated with a black line and taxonomy. **b.** Genome recovery metrics for putative methanotrophs. The number of genomes recovered, novel species recovered, number of high quality (HQ) genomes recovered and number of HQ genomes of novel species are displayed. The values are square root transformed and indicated by separate patterns. Shown are genera containing at least one novel species of putative methanotrophs.

**Figure 2:**
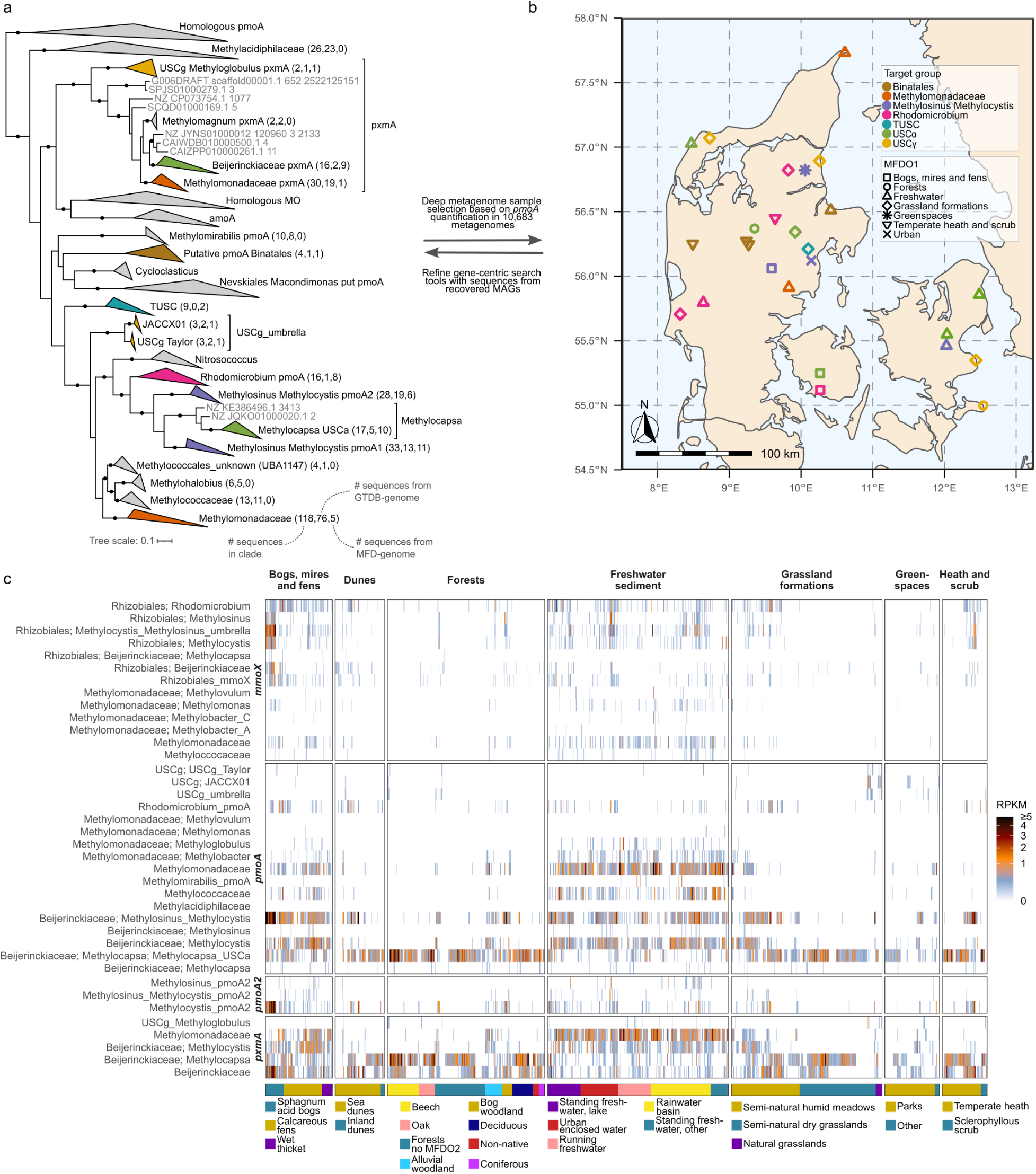
Quantification of methanotroph marker genes in Danish natural habitats. **a.** Protein phylogenetic tree of PmoA sequences. Clades of sequences targeted by long read metagenomic sequencing are coloured and correspond to samples selected in b). Numbers indicate the total number of sequences in a given clade, the number of sequences obtained from GTDB r220 genomes, and the number of sequences obtained from the genomes recovered here. Bootstraps with above 95% support (UFB, 1000 replications) are indicated by a black circle. **b.** Map of Denmark with the 26 selected samples for long read sequencing displayed. The samples are coloured by the target clade, and the shape indicates the MFDO1 habitat type. **c.** Distribution of methanotrophy marker genes in major natural habitats in Denmark. Samples are coloured by the number of reads (Reads Per Kilobase Million (RPKM)) assigned to each gene-phylogenetic group. The gene-clades correspond to clades in the protein phylogenetic tree in a. Samples are clustered with hierarchical clusters within each MFDO2 habitat. The bottom colour panel indicates the MFDO2 habitat.

### Diversity and habitat preferences of Danish methanotrophs

To determine the distribution of Danish methanotrophs, we applied our tailored search model to the 10,683 Microflora Danica metagenomes^2^. The metagenome samples are accompanied by a detailed description of sample type (i.e. soil or sediment), habitat type (i.e. natural, agricultural or urban) and three-level habitat descriptions (i.e. MFD ontology level 1 (MFDO1): grassland formations, MFDO2: semi-natural humid meadows, MFDO3: molinia meadows). We show that all sampled natural soils and sediments contained methanotrophs, with distinct differences in the methanotrophic community between habitats (**Figure 2**).

The highest relative marker gene abundances *(pmoA/mmoX)* of canonical methanotrophs were found in the bogs, mires and fens (MFDO1) and freshwater sediments (MFDO1) (**Figure 2c**). Within the bogs, mires and fens, the sphagnum acidic bogs (MFDO2) were characterised by a high relative abundance of *Methylosinus/Methylocystis mmoX* and *pmoA*, along with the associated paralogs *Methylosinus/Methylocystis pmoA2* and *pxmA*. *Methylocystis/Methylosinus pmoA* and *pxmA* were also abundant in the calcareous fens (MFDO2) and wet thickets (MFDO2), though less so than in sphagnum acidic bogs (**Figure 2c**). *Methylomonadaceae pmoA* and *pxmA* (e.g. *Methylobacter, Methyloglobulus*) were generally more prevalent in calcareous fens and wet thickets than in acidic bogs. This supports previous work on the preferences of *Methylocystaceae* methanotrophs for low nutrient concentrations and pH in bogs in contrast to the preferences of *Methylomonadaceae* methanotrophs for higher nutrient concentrations and pH in fens^12,48,49^. Freshwater sediments contained high diversity and relative abundance of *Methylocystis/Methylosinus pmoA* and *mmoX* and *Methylomonadaceae pmoA, mmoX* and *pxmA*. Additionally, *Methylococcaceae pmoA* (e.g. *Methylococcus*, *Methylocaldum, Methylomidiphilus*) were also present in 26% of bogs, mires and fens and 36% of freshwater sediments (**Figure 2c**). Sub-habitats of dunes, forests, meadows and heaths that are either permanently or periodically wet include humid dune slacks (MFDO3) and decalcified Empetrum dunes (MFDO3), alluvial-and bog woodland (MFDO2), hydrophilous tall-herb swamp (MFDO3), molinia meadows (MFDO3) and wet heath (MFDO3). In these habitats, we also observe *Methylocystis/Methylosinus pmoA* and *mmoX,* with *Methylomonadaceae pmoA* and *pxmA* occurring in the hydrophilous tall-herb swamps (**Figure 2c, Extended Data Figure 4**).

Genes associated with atmospheric methane consumption in upland soils include those assigned to USCα and USCγ. Among these, *Methylocapsa-*associated USCα *pmoA* were found in nearly all sampled habitats, while USCγ *pmoA* were only present in a few greenspaces and grasslands (**Figure 2c**). *Methylocapsa* USCα *pmoA* were abundant in the dry subhabitats within dunes, forests, grasslands, greenspaces and heath and scrub (**Figure 2c, Extended Data Figure 4**), often occurring with *Methylocapsa pxmA* as the only methanotrophy genes in these habitats. In urban greenspaces (**Figure 2c**) and agricultural fields (**Extended Data Figure 4**), *Methylocapsa* USCα *pmoA* and *Methylocapsa pxmA* were also the most prevalent genes, but were detected inconsistently, independent of crop type and in lower relative abundance (greenspaces: Mann–Whitney U-test, U = 1,117,357, two-sided, P = 8.47 × 10^−98^, n=708); agriculture: Mann–Whitney U-test, U = 1,651,720, two-sided, P = 9.69 × 10^−197^, n=2945) compared to natural upland soils (semi-natural dry grasslands (MFDO2), dry heaths (MFDO3) and forests types beech, oak, deciduous, coniferous and non-native trees (MFDO2), n=2094). This aligns with recent findings of Danish agricultural soils as minor methane sinks on average, with uptake varying markedly across locations, influenced by soil-structure and moisture rather than crop type^50^. While organic fertilisers have been added to induce transient methanotrophic methane oxidation in agricultural upland soils^51–53^, the origin of this enhanced methane uptake is unclear. Proposed mechanisms include the introduction of methanotrophs (and methanogens) via the added compost^53^, or activation of native canonical MOB^51,52,54^. We observed no canonical MOB in most Danish agricultural soils (**Extended Data Figure 4**), and generally observed lower MOB diversity and relative abundance in highly managed habitats. Consistent with the sensitivity of the atmospheric methane sink to disturbances^55–57^, our work supports that the upland soil methane sink is impaired by the conversion to agriculture.

### *Methylocapsa* are widespread in natural upland soils

Several of the most widespread and abundant methanotrophy marker genes were associated with newly recovered genomes (**Figure 1, 2**). To assess the potential roles of the corresponding organisms as methane sinks, we connected their genomic potential to specific habitat niches by genome-level quantification using sylph^58^. Novel and known members of *Methylocapsa* were amongst the most abundant and widespread, and comprised a third (33%) of the new methanotroph species recovered (**Figure 1 and 2**). The majority of the recovered *Methylocapsa* species (27) encoded the *Methylocapsa* USCα *pmoA*, but some encoded canonical forms of *pmoA* and *mmoX* instead (**Figure 3a**). We evaluated methane associated metabolism by identifying carbon assimilation-and dissimilation modules for methanol, formaldehyde and formate. *Methylocapsa* members encoded the MDH-*xoxF* for methanol oxidation, the H_4_MPT pathway for formaldehyde oxidation, the H_4_F pathway for formate assimilation and the serine cycle for carbon assimilation (**Figure 3a, Extended Data Figure 5**). Beyond these similarities in carbon assimilation and dissimilation, *Methylocapsa* clades displayed distinct habitat preferences and differences in metabolic potential.

**Figure 3:**
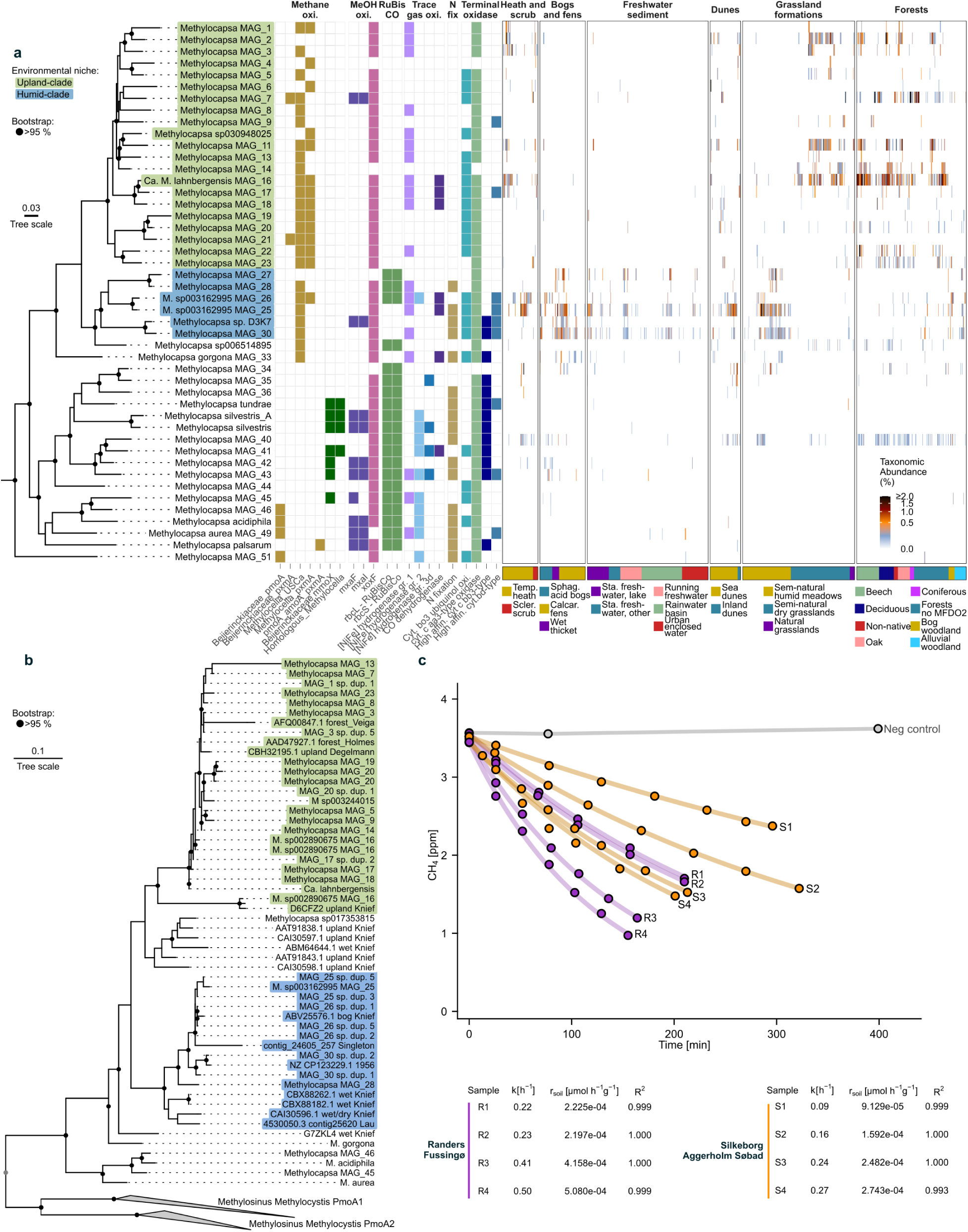
Quantification and characterisation of *Methylocapsa* MAGs. **a.** Phylogenomic tree of *Methylocapsa* members from GTDB and MFD recovered MAGs. Genomes have been dereplicated at 95% ANI. Bootstraps with above 95% support (UFB, 1000 replications) are indicated by a black circle. Two distinct clades are indicated in green (upland clade) and blue (humid clade). Associated with each genome is the phylogeny of *pmoA, pxmA* and *mmoX,* along with metabolic potential for methanol oxidation, CO_2_-fixation, trace gas oxidation (H_2_ and CO), nitrogen fixation and type of terminal oxidase. Each genome is quantified using sylph, with taxonomic abundance shown across the major natural habitats. Habitats are clustered (hclust) within MFDO2. **b.** Protein phylogenetic tree of *Methylocapsa* PmoA sequences and USCα PmoA sequences from amplicon studies as reported in^12^. Clades of PmoA sequences are coloured corresponding to the genomes from which the PmoA sequence was recovered. Bootstraps with above 95% support (UFB, 1000 replications) are indicated by a black circle. The tree was generated with all sequences in the *pmoA* GraftM package with the protein model WAG, and subsequently subsetted to PmoA sequences from *Methylocapsa, Methylocystis* and *Methylosinus*. **c.** Oxidation rate experiment of selected soils from Randers, Fussingø (Beech forest) and Silkeborg, Aggerholm Søbad (Deciduous forest). Headspace concentrations of methane were measured from four replicates of each soil. Negative control was made with autoclaved soil from Silkeborg. A linear model (function lm()) of the natural log of methane concentrations was calculated to find first order decay (k, h^-1^) constant and correlation coefficient.

A monophyletic clade consisting almost exclusively (21/23) of our newly recovered *Methylocapsa* MAG spp. displayed high relative abundance in upland soils including forests, dry grasslands and temperate heaths (**Figure 3a**, light green clade, now referred to as “upland clade”). This clade encoded the *Methylocapsa* USCα *pmoA* similar to those frequently identified in upland soils^12^ and the *Methylocapsa pxmA,* with some variability between species likely due to genome incompleteness (**Figure 3a,b**, upland clade, **Extended Data Figure 5**). Abundant species of these potential atmMOB include *Methylocapsa* MAG_2, MAG_7 and MAG_11. Intriguingly, *Methylocapsa* MAG_7 was highly abundant in some beech, oak, deciduous and coniferous forests as the only member of *Methylocapsa*, and was nearly absent in all other samples. The species cluster of “*Methylocapsa* MAG_7” consisted of 23 MAGs with completeness >80% and contamination <10%, of which 20 encoded the calcium-dependent *mxaF*-MDH in addition to xoxF-MDH (**Extended Data Figure 5**). Other novel *Methylocapsa* species found in upland soils (MAGs 16, 17, 18, upland clade) formed a monophyletic clade with *‘Candidatus* Methyloaffinis lahnbergensis*’* (part of *Methylocapsa* using GTDB-taxonomy), a proposed atmMOB showing active pMMOs at atmospheric methane levels^59^. *Ca.* M. lahnbergensis and associated species were particularly abundant in dry grasslands and across most forests in Denmark, and showed genomic potential for H_2_ (NiFe gr. 1) and CO oxidation (**Figure 3a, Extended Data Figure 6**). The genomic potential of the *Ca.* M. lahnbergensis associated clade to obtain energy from several trace gases could explain its prevalence in upland soils.

A second *Methylocapsa* clade (**Figure 3a**, blue clade, now referred to as “humid clade”) encoded *Methylocapsa* USCα *pmoA* similar to those identified in wet soils (**Figure 3b**). This clade included *Methylocapsa* D3K7 and a MQ MAG sp003162995 (recovered from Stordalen Mire), and was abundant in calcareous fens, certain sediments and sea dunes, alluvial-and bog woodlands and humid meadows. We observed strong habitat partitioning between the two *Methylocapsa* clades (upland-vs humid clade), with mutual exclusion in heath and shrub (J=0.0579, p=5.46e-27), dunes (J=0.0876, p=7.86e-3) grasslands (J=0.0519, p=5.96e-28) and forests (J=0.0670, p=2.31e-13) (**Figure 3a**). Habitats preferred by the humid clade are permanently or periodically wet and anticipated to exhibit conditions of oxygen limitation. While these members also encode the *Methylocapsa* USCα *pmoA*, most also hold the genomic potential to fix CO_2_ through the CBB cycle, oxidise H_2_ and CO, fix N_2_ and use sub-atmospheric levels of oxygen (cytochrome bd ubiquinol oxidase) (**Figure 3a, Supplementary Note 2**).

Additionally, *pxmA* was encoded in most *Methylocapsa* upland-clade species, and was absent in most *Methylocapsa* in the humid clade (**Figure 3a**). Although the function of pXMO is unknown, *pxmABC* is expressed in response to O_2_-limited conditions in the Gammaproteobateria *Methylomonas denitrificans* FJG1^47^ and *Methylomicrobium album* BG8^60^. In contrast, the gene and genome-based analyses of the *Methylocapsa* MFD putative MOB indicate the *pxm-*containing groups do not occupy wet environments, and instead occur in upland soils where atmospheric methane uptake is suspected.

While the USCα form of pMMO is associated with atmMOB, we show here that the accompanying metabolic potential markedly affects the niches of individual *Methylocapsa* clades. The *Methylocapsa* encoding canonical *pmoA* or *mmoX* did not appear to be abundant in Danish natural habitats (e.g. *M. silvestris, M. acidiphila, M. aurea, M. palsarum,* all *Methylocapsa,* following GTDB taxonomy). Likewise, the atmMOB *M. gorgona* did not appear abundant, with genome-and PmoA phylogeny distinct from the novel and abundant potential atmMOB recovered in this work (**Figure 3a,b**).

From our analysis, it appears that the Danish sink for atmospheric methane is dominated by few (10-15) *Methylocapsa* spp., many of which were identified first here (**Figure 3a**). As a demonstration of the ecophysiological insight offered by the genome-resolved approach, we wanted to quantify the potential of *Methylocapsa* MAG_7 to consume atmospheric methane, due to its high taxonomic abundance (sylph) in upland forest soils (**Figure 3a,c**). To monitor methane oxidation rates over time, soil samples from two forests dominated by *Methylocapsa* MAG_7 were incubated in airtight glass bottles with methane concentrations slightly above atmospheric levels (∼3.5 ppm_v_). Methane was oxidised to below atmospheric concentrations within hours, with the decline in concentration following first order kinetics (Randers, Fussingø: k=0.2-0.5 h^-1^, R^2^>0.998, r_soil_=2.2-5.1×10^-4^ µmol h^-1^ g_soil_^-1^ at 15°C, 1.013 bar, 1.93 ppm CH_4_; Silkeborg, Aggerholm Søbad: (k=0.09-0.27 h^-1^, R^2^>0.993, r_soil_=0.91-2.7×10^-4^ µmol h^-1^ g_soil_^-1^ at 15°C, 1.013 bar, 1.93 ppm CH_4_) (**Figure 3c**). These oxidation rates are comparable to those reported from Rold Skov, Denmark^8^, and noticeably higher (∼10x) than those reported in German forests^59^. Following the experiment, we used deep long-read sequencing to confirm the presence of *Methylocapsa* MAG_7. Both soil samples yielded circular HQ genomes of the same species (>99% ANI to *Methylocapsa* MAG_7, 1% relative abundance), and the only methanotrophic species recovered were two other new *Methylocapsa* spp. (0.4% relative abundance, HQ MAG; 0.06%, MQ MAG) (**Supplementary Data File 1**). By uncovering the diversity, distribution and metabolic potential of putative Danish methanotrophs in soils with methane sink potential, our genome-resolved approach identifies the key putative atmMOBs for directed future ecophysiology studies.

### Methylocystis species are ubiquitous in wetlands and sediments

We recovered 20 new species within the genus *Methylocystis*. While this genus is well described using isolates, through our new species we identified sub-populations with distinct niches and differences in habitat prevalence that were linked to *pmoA* and *pmoA2* phylogeny. Members of *Methylocystis* encoded the pMMO, and occasionally sMMO (**Figure 4a**). Furthermore, nearly all members encoded at least one type of hydrogen dehydrogenase and the ability to fix nitrogen (**Figure 4a**). One clade (**Figure 4a**, light blue clade, now referred to as “clade 1 (fen)”) included *M. silviterrae* and *M. rosea*, and members were very abundant in calcareous fens and wet thickets, alluvial woodland, nearly all freshwater sediments and most humid meadows. These *Methylocystis* spp. encoded distinct clades of *pmoA1* and *pmoA2,* and a copy of the *pxmA* (**Figure 4a**). Furthermore, most members had an *mxaF*-type methanol dehydrogenase in addition to the *xoxF*-type, and a high-affinity cytochrome bd in addition to caa_3_-type. While *M. rosea* can grow on air in flowing filter cultures^27^, we did not observe *M. rosea*, nor the associated clade of *pmoA1,* to be present in upland soils where air is considered the main source of methane (**Figure 4a**, **Figure 2c**). Beyond this, *M. silviterrae* and *M. hirsuta* also encoded the sMMO (*mmoX*), and appeared exclusively abundant in a subset of urban enclosed water samples on Zealand (**Figure 4b**).

**Figure 4:**
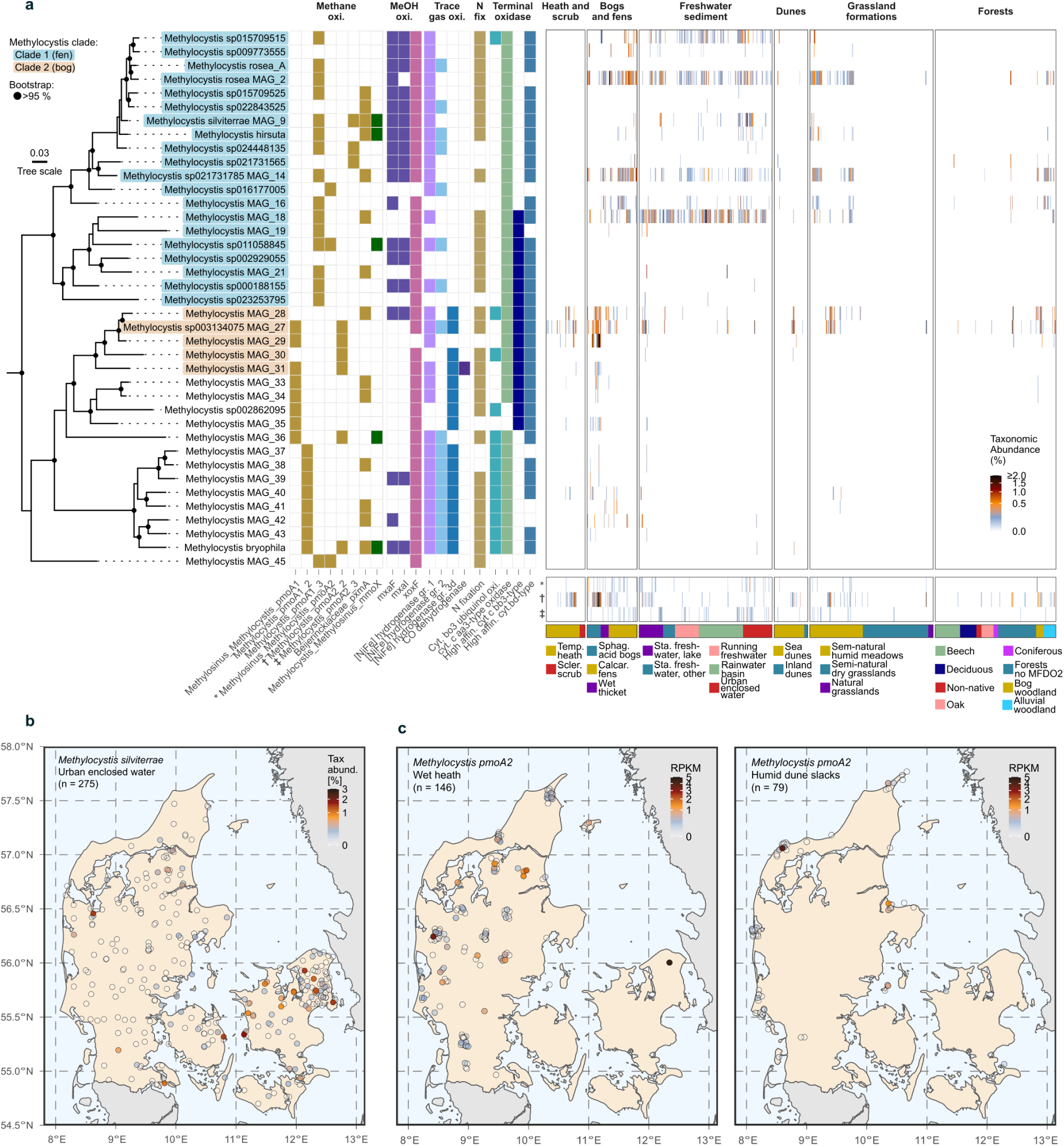
Quantification and characterisation of *Methylocystis* MAGs. **a.** Phylogenomic tree of *Methylocystis* members from GTDB and MFD recovered MAGs. Genomes have been dereplicated at 95% ANI. Bootstraps with above 95% support (UFB, 1000 replications) are indicated by a black circle. Associated with each genome is the phylogeny of *pmoA, pmoA2* and *mmoX,* along with metabolic potential for methanol oxidation, trace gas oxidation (H_2_ and CO), nitrogen fixation and type of terminal oxidase encoded in the electron transport chain. Additionally, each genome has been quantified (sylph), and the taxonomic abundance is shown across the major natural habitats. Below, the relative read count (RPKM) of the genes (*) ‘Methylosinus_Methylocystis_pmoA2’, (*†*) ‘Methylocystis_pmoA2_2’ and (*‡*) ‘Methylocystis_pmoA2_3’ are displayed, corresponding to the *pmoA2* genes encoded in *Methylocystis* genomes. Habitats are clustered (hclust) within MFDO2. **b.** Taxonomic abundance (sylph) of *M. silviterrae* MAGs within the MFDO2 habitat Urban enclosed waters. The location of all samples within the Urban enclosed water are shown in Denmark (map from EuroGeographics). **c.** Quantification of *Methylocystis pmoA2* gene abundance (RPKM) in the MFDO3 habitats wet heath and humid dune slack (map from EuroGeographics). All samples within the wet heath and humid dune slacks are displayed on the map.

A second population of *Methylocystis* spp. (**Figure 4a**, beige clade, now referred to as “clade 2 (bog)”) encoded distinct clades of *pmoA1* and *pmoA2,* as well as an additional hydrogenase (NiFe group 3d), and a high-affinity cbb_3_-type cytochrome c oxidase in addition to caa_3_ type and cytochrome bd (**Extended Data Figure 6**). The clade 2 (bog) had no isolate representatives but contained a MAG recovered from Stordalen Mire (*Methylocystis* sp003134075). Between the two abundant clades (clade 1 (fen), and clade 2 (bog)) we observed contrasting habitat preferences, likely linked to pH and nutrient availability, with mutual exclusion in acidic bogs and calcareous fens (bog, mire and fens: J=0.107, p=7.63e-29), bog-and alluvial woodlands (forests: J=0.0382, p=8.48e-9) and molinia meadow and hydrophilous tall-herb swamp (humid meadows: J=0.0606, p=9.08e-7) (**Figure 4a**).

We found the highest relative gene abundance of *Methylocystis pmoA2* and taxonomic abundance of the corresponding *Methylocystis* sub-population (clade 2 (bog)) in wet habitats, namely in sphagnum acid bogs and bog woodland, and a subset of the MFDO3 habitats humid dune slacks, decalcified Empetrum dunes, molinia meadows and wet heath (**Figure 4a, Extended Data Figure 4**). This was surprising, as *pmoA2* transcripts in *Methylocystis* SC2 have been linked to atmospheric methane oxidation in culture^26^. Additionally, the relative abundances of *Methylocystis pmoA2* varied greatly even within nearby samples from the same MFDO3 habitat (e.g. wet heath and humid dune slacks) (**Figure 4c**). While transcription of *pmoA2* has been linked to methane oxidation at low concentrations in rice paddy fields^61^ and in drained peatland grasslands^62^, these are both habitats with seasonal methane production, and might be explained by the temporary ability of *Methylocystis* spp. to oxidise atmospheric methane after exposure to high levels of methane^13,25^. Here, we observe no pattern in the distribution that might suggest *Methylocystis,* with or without *pmoA2,* to inhabit sites with consistent limitation in methane supply. Instead, the abundant members in Danish forests, dry heaths, meadows and grasslands, where air is the main source of methane, are MOB associated with *Methylocapsa* encoding USCα *pmoA*.

### Gammaproteobacterial methanotrophs have limited distributions

Gammaproteobacterial methanotrophs, from which we recovered 32 new species (**Figure 2**), differ in adaptation to temperature, pH, salinity and oxygen levels, but are often identified in aquatic habitats^12^. Putative methanotrophs were recovered in the genera *Methylobacter, Methylobacter_A, Methylobacter_C, Methylomonas, Methyloglobulus, Methylovulum,* UBA10906 and JAGXGJ01, all encoding the RuMP pathway for formaldehyde assimilation and additional H_4_MPT pathway for formaldehyde oxidation (**Extended Data Figure 7a, Extended Data Figure 8a**). Across Denmark, we found MAGs classified as *Methylobacter_C* highly abundant in freshwater sediments (**Extended Data Figure 7a**). Several species (4/9), including the most widespread and abundant *Methylobacter_C* sp002256465 MAG_17, encoded the potential for partial denitrification to nitrous oxide (*narGHJI, nirK, norCB*) (**Extended Data Figure 8b**). Coupling of methane oxidation with nitrate respiration has been shown in members of *Methylomonas denitrificans*^47^ and populations of *Methylomonas* and *Methylobacter* have been proposed to oxidise methane using nitrate reduction^63^.

Putative gammaproteobacterial atmMOB encoding USCγ are suggested to undertake atmospheric methane oxidation in soils with pH>6^14,24,30,64^. We recovered novel USCγ-encoding genomes from the genera USCg-Taylor (1 HQ, 1 MQ) and JACCXJ01 (1 MQ). As with genomes already deposited in GTDB r220 (**Extended Data Figure 2**) and ‘*Ca.* Methyloligotropha calcicola’ MAGs recovered from limestone caves^31^, methanol oxidation was encoded by MDH-*xoxF*, and formaldehyde oxidation through the H_4_MPT pathway, but the key genes in the RuMP (*HxlA/hps* and *HxlB/phi*) and serine cycle (*hprA*, *gck*) were not identified. Two USCg-Taylor genomes also encoded a copy of the *pxmA* (**Extended Data Figure 7a**). The USCγ-affiliated genomes, USCγ *pmoA* and USCγ-related *pxmA* genes were predominantly restricted to the grassland habitats calcareous grassland (MFDO3) and xeric sand calcareous grasslands (MFDO3) and a few greenspaces (**Extended Data Figure 7a, Extended Data Figure 4a**). Furthermore, the USCγ *pmoA* and *Methylocapsa* USCα *pmoA* were not present in the same grassland formations (J=0.0199, p=0.0000860).

To investigate the methane oxidation kinetics from soils containing the USCγ *pmoA* encoding putative atmMOB, we resequenced promising grasslands with a high taxonomic abundance of JACCXJ01 MAG_1, and selected a xeric sand calcareous grassland for methane oxidation rate experiments as described above. We observed oxidation of methane to below atmospheric concentrations following first order kinetics in two (0.097-0.122 h^-1^, R^2^>0.998, r_soil_=0.97-1.2×10^-4^ µmol h^-1^ g_soil_^-1^ at 15°C, 1.013 bar, 1.93 ppm CH_4_) of the four replicates from the site (**Extended Data Figure 7c**). The lack of activity in two of the samples could reflect the spatial heterogeneity of USCγ, which was also indicated by its varied detection in co-located samples (**Extended Data Figure 7b**). From the oxidation rate samples, we used deep long-read sequencing to assemble three circular HQ JACCXJ01 MAGs of three new species (<95% ANI), with relative abundances of 0.25, 0.4 and 0.6% (**Supplementary Data File 1**). Additionally, one HQ MAG of a new USCg-Taylor species was recovered at a relative abundance of 0.12%. No other canonical or putative methanotrophs were assembled from the soil. With our genome-resolved approach, we identified putative USCy atmMOB and demonstrate their potential involvement in methane oxidation and contribution to methane sink functionality of calcareous soil types that should be examined in the future.

### Putative methanotrophs from low abundance populations display unique metabolisms

The recent confirmation of methanotrophy and thiotrophy within a single organism, *Methylovirgula thiovorans*^46^, isolated from a Korean peatland, prompted us to investigate this metabolism in Danish bogs and fens. We recovered 15 MAGs (6 novel species) within the genus *Rhodomicrobium* encoding putative PmoA and MmoX similar to the “HYP” PmoA and MmoX sequences^22^ (78-98% and 62-99% aa identity, respectively) and to the “Novel” PmoA sequences^65^ (84-87% aa identity). Putative *Rhodomicrobium* PmoA and MmoX sequences formed distinct monophyletic clades in protein trees (**Figure 2a**), and the *Rhodomicrobium* MAGs encoding MMOs clustered together in a phylogenomic tree (**Extended Data Figure 10a**). Downstream metabolic repertoire for methane oxidation was encoded by a MDH-*xoxF*, GSH-linked formaldehyde oxidation and the CBB cycle for carbon assimilation, along with formate oxidation to CO_2_ (**Extended Data Figure 10a**). Additionally, the putative methanotrophic *Rhodomicrobium* spp. also held the potential to oxidise reduced sulphur compounds, encoding sulfate adenylyltransferase (*sat*), adenylyl-sulfate reductase (*aprAB)* and reverse dissimilatory sulfite reductase (*rdsrAB,dsrC,dsrEFH*), confirmed by protein phylogeny (**Extended Data Figure 10b**), with the remaining operon following the expected structure for sulfur-oxidisers^66^. Our putative methanotrophic *Rhodomicrobium* MAGs also encoded a high affinity cbb_3_-type cytochrome c oxidase, potential to fix nitrogen (*nifHDK*), oxidise carbon monoxide (*coxSML*) and hydrogen ([NiFe] gr. 1, 2 and 3d) and the potential to store poly-3-hydroxybutyrate (PHB) (**Extended Data Figure 10b**), similar to the metabolic potential of *Rhodomicrobium* sp. HYP1^22^ and *M. thiovorans*^46^. The proposed novel group of methanotrophs within *Rhodomicrobium,* and corresponding *Rhodomicrobium pmoA* and *mmoX* genes, could be identified in similar habitats as the *Methylocystis/Methylosinus* clade (e.g. calcareous fens, humid dune stacks, wet heath), albeit in lower relative abundance compared to canonical MOB genera (**Figure 2, Extended Data Figure 10a**). Uncovering the potential role of combined methano-and thiotrophy is of interest given the tight linkage of methane and sulphur cycles in peatland and sediments^67,68^.

Efforts to identify *pmoA*-like sequences in upland soils have led to the hypothesis that the TUSC/Cluster-2 group is linked to atmospheric methane consumption^69–71^. To investigate this, we linked this ambiguous clade to several new genomes within a single genus of the class Binatia and explored their methanotrophic potential. We recovered seven MAGs from five novel species containing TUSC-like *pmoA* sequences and 81 MAGs from 15 novel species encoding Binatales *pmoA-*like sequences (**Extended Data Figure 11a**). All members belonged to Binatia, within the orders JACPRU01, Binatales or Bin18. While some (3/7) JACPRU01 MAGs encoded a complete GSH-linked pathway for formaldehyde oxidation, the remaining modules for methanol oxidation, and formaldehyde or formate assimilation were incomplete or missing (**Extended Data Figure 11a**). We identified TUSC genes in the genus JACPRU01 similar to original TUSC-sequences (86-100% aa identity, tblastn against AJ579663, AJ868246, EU723743, KC122308)^69–71^. The “Putative *pmoA* Binatales” and TUSC clades did not cluster with verified *pmoA* sequences, but are located in a part of the tree with hydrocarbon monooxygenases (**Figure 2**)^72,73^. Alternative metabolisms based on the associated genomes included denitrification (*narGH*, *nirK*, *norCB* and *nosZ)* (**Extended Data Figure 12**). Despite the lack of genomic evidence for methane oxidation, TUSC associated sequences were identified in a subset of fens, fields, forests, grasslands and greenspaces, with a distribution pattern including but not limited to upland soils (**Extended Data Figure 11b**). The apparent lack of central methane-processing metabolism highlights the importance of a genome-based analysis involving phylogeny and assessment of multiple metabolic modules, preventing misinterpretation if analyses are based on single genes or genomes.

## Conclusions

Here we provide the first country-wide description of aerobic methanotrophs, their habitat preferences and biogeography. By applying large-scale, genome-resolved metagenomics, we reveal new, abundant and functionally relevant species that could not be resolved using gene-based approaches. Uncharacterised *Methylocapsa* spp. show strong niche partitioning in natural upland soils and represent potentially important methane sinks. In contrast to natural soils, methanotrophs were mostly absent from habitats with human interference, underlining the impact of disturbance on soil methane sink potential. In wet environments, undescribed species from *Methylocystis* and *Methylobacter_C* were abundant. Their role as methane filters will be important to monitor, especially following “The Agreement on a Green Denmark” as large areas of drained agricultural land will be restored. Furthermore, we show the importance of evaluating new species in the context of genomic potential and ecological niche. This is demonstrated by the metabolic potential of methano-and thiotrophy within *Rhodomicrobium*, the lack of methanotrophic metabolic potential in Binatia MAGs encoding TUSC sequences, and the habitat specific and phylogenetic signal of *Methylocapsa* encoding USCα *pmoA* in humid versus dry environments. Our results provide a foundation for ecosystem modelling based on the physiology of locally abundant and important taxa, and highlight the need for high resolution studies in other biogeographic regions to determine the relevance of Danish methanotroph spp. to global distributions.

## Data availability

Methanotroph genomes from the Microflora Danica long-read MAG catalogue are available at ENA with BioProject ID: PRJEB58634. Individual accession numbers for methanotroph genomes used in the study are provided in (**Supplementary Data File 2**). Moreover, methanotroph genomes from the Microflora Danica short-read MAG catalogue are available at the NCBI GenBank under BioProject PRJNA1071982. Sequencing data and MAGs recovered from soil samples post oxidation rate experiments are available at ENA with BioProject ID: PRJEB104930. Public data used in this study include GTDB r220 (https://data.gtdb.ecogenomic.org/releases/) and KEGG database release 109 (https://www.kegg.jp/kegg/docs/relnote.html). GraftM packages *mmoX* and *pmoA* updated with sequences recovered in this study can be found at GitHub (https://github.com/KalinkaKnudsen/MFD_methanotrophs_DK).

## Acknowledgements

We thank the Microflora Danica Consortium for their contributions to sample and metadata collection across Denmark; M Chuvochina for her help in checking proposed names for select microbial lineages. Funding was provided by the Poul Due Jensen Foundation, PDJF (grant MicroFlora Danica to M.A. and P.H.N.), Villum Foundation (grant 15510 and 50093 to M.A., grant 13351 to P.H.N., grant VIL60768 to C.M.S), the European Union (ERC grant 101078234 to M.A.).

C.M.S. was supported by a Novo Nordisk Foundation Postdoctoral Fellowship grant (NNF20OC0065005), and F.D. was supported by a Novo Nordisk Foundation Challenge Programme grant (NNF24OC0085294). A.T.T. was supported by a Research Council of Norway Young Researcher grant (project Living on air 315129).

## Author Contributions

C.M.S., K.S.K., M.A., P.H.N. designed the study. M.S., C.J., R.H.K. and K.S.K. performed DNA extraction and processing of samples for Nanopore sequencing. K.S.K. selected samples for long-read sequencing. K.S.K. and F.D. conducted resampling for oxidation rate experiments. M.S. performed MAG recovery. K.S.K., T.S. and A.T.T. performed and designed oxidation rate experiments. A.T.T. and T.S. provided input and specialist knowledge. F.D. and T.B.N.J. performed curation and validation of the sample metadata. K.S.K. constructed marker gene databases, analysed metagenomes, and conducted metabolic annotation, phylogenetic analysis, and wrote the initial manuscript. K.S.K., C.M.S., M.A. wrote the manuscript, with input from all authors. All of the authors reviewed and agreed with the manuscript.

## Methods

### Curation and generation of HMM search packages

HMM search packages were generated by using the *pmoA* and *mmoX* GraftM packages from Singleton et al. 2018. The packages were amended with sequences produced in recent studies^12,34–46^. The updated packages were used to screen GTDB r220. The potential hits obtained were filtered based on sequence phylogeny and length of translated protein (minimum 240aa PmoA and 400 MmoX) and HMM alignment score < 1e-10. Hits were translated to protein sequences using Prodigal^74^. Sequences were dereplicated using usearch^75^ v11.0.667 option – cluster_fast. Groups were differentially dereplicated to not bias the HMM towards sequences clades with many representatives. Thus, *Methylococcaceae pmoA* and *pxmA* sequences were dereplicated at 95% aa. identity. Remaining *pmoA* sequences were clustered at 99% aa identity. All *mmoX* sequences were clustered at 99% aa identity. Multiple sequence alignment was performed using MAFFT^76^ v7.490. Trimming was performed at minimal 20% representation at a given site using trimAl^77^ v1.5.0. Protein phylogenetic trees were generated with IQ-TREE^78^ v. 2.1.2, with the ultrafast bootstrap approximation option using 1000 iterations, and enabling the ModelFinder option (best fit model pmoA: LG+F+R7, mmoX: LG+R5). The trees were rerooted and grouped in ARB^79^ v. 6.0.3. Visualisation of protein trees was done in iTOL^80^ v6. Protein trees were manually grouped and rerooted before submission to GraftM. Groups were inferred based on gene-phylogeny from^22^ and taxonomy of the genomes from which sequences were obtained in GTDB r220. General search and manipulation of protein and nucleotide sequence files was performed using SeqKit^81^ v2.0.0. HMM search packages were created using GraftM^21^ v0.15.0. Contamination was evaluated by searching with blastp against nr (**Supplementary Note 1**). Additionally, GUNC^82^ v1.0.6 was used for extra contamination evaluation for the Ca. M. Kingii MAG, with the database gunc_db_progenomes2.1. Finally, potential methanotroph MAGs recovered in^3^ and^2^ were identified by running the updated GraftM packages (*pmoA, mmoX*) on all MAGs recovered. Potential hits were evaluated based on phylogenetic placement and hits were filtered with a conditional E-value (c-Evalue) threshold of 1e-10. These sequences were dereplicated at 100% identity and added to construct the final GraftM *pmoA* and *mmoX* packages. The GraftM packages are available at GitHub (https://github.com/KalinkaKnudsen/MFD_methanotrophs_DK).

### Gene abundances across samples

Short-read samples from Microflora Danica^2^ were filtered to include only samples with at least 500,000 reads. Then, GraftM^21^ was used to quantify *pmoA* and *mmoX*-like sequences in the samples, using hmmsearch+diamond search methods. Hits were filtered with a conditional E-value (c-Evalue) threshold of 1e-10. Parallelisation was applied when possible using GNU parallel^83^. Read abundance was normalised by HMM-alignment length (pmoA: 927nt, mmoX: 1581nt) and reported in reads per kilobase million (RPKM). Processing and normalisation was carried out in R^84^ v4.4.0. The main displays show only gene-phylogenetic clades of interest, and excludes hits classified as hydrocarbon or ammonia monooxygenases. All hits can be found in (**Supplementary Data File 4**). Samples were clustered within the MFDO2 habitats based on the hellinger-transformed Bray-Curtis dissimilarity matrix (function vegdist, method=”hellinger”, package vegan^85^ v2.6.6.1). Clustering was carried out with the stats^84^ v4.4.1 package (function *hclust,* method=ward.D2). The MFDO2 habitat for “Soil, Natural, Forests” was replaced with the MFDO3 habitat, since the MFDO2 was not descriptive for the analysis performed here, being either “Temperate forests” or “Forest (non-habitattype)”

### Genome abundance

Quantification of genomes within the shallow metagenomes was carried out using sylph^58^ v0.6.1. The genome catalogue was composed of dereplicated MAGs recovered in Microflora Danica^2,3^, along with GTDB r220. In cases where a species representative was present in both GTDB r220 and the Microflora Danica genome catalogue, both were quantified and subsequently aggregated in R^84^ v4.4.0 with tidyverse^86^ v2.0.0, as we wanted a local representative but did not want to exclude genomes from the GTDB r220 that might be divergent. The genome catalogue and samples were independently prepared by the “sketch” command and default parameters. Then, the taxonomic abundance of the genomes in the samples was determined using the “profile” command. Taxonomy was added using the utility script sylph_to_taxprof.py, and the quantification was extracted using the utility script merge_sylph_taxprof.py.

### Co-occurance of genes and genomes

Abundance profiles of genes or genomes within each MFDO1 did not follow normal distribution (determined using the shapiro test with rstatix^87^ v0.7.2 package). Therefore, a non-parametric approach was applied in all downstream analysis. Between groups of genes or genomes, the Jaccard similarity was calculated based on presence and absence gene abundances with the formula J(geneA, geneB)=|geneA ⋂ geneB|/(|geneA ⋃ geneB) in R^84^ v4.4.0. To accompany the Jaccard similarity index, a Fisher’s Exact Test was conducted using the function *fisher.test()* (stats^84^ v4.4.0). P.values were evaluated as one-sided “less” in cases where genes were mutually exclusive, and as one-sided “greater” in cases where genes co-occured, and were adjusted for multiple comparisons using the Benjamini-Hochberg (BH) approach (function p-adjust(), stats^84^ v4.4.0. Maps displaying abundance of genes or genomes were produced in R^84^ v4.4.0 with packages sf^88^ v1.0.21 ggspatial^89^ v1.1.10 and ggtext^90^ v0.1.2. The background map was downloaded from EuroGeographics.

### Identification of Methanotroph MAGs and annotation

Potential methanotroph MAGs were identified by running the updated GraftM packages (*pmoA, mmoX*) on all MAGs recovered in^3^ and^2^. Potential hits were evaluated based on phylogenetic placement and hits were filtered with a conditional E-value (c-Evalue) threshold of 1e-10. Subsequent annotation of MAGs was performed with DRAM^91^ v1.4.6 and the KEGG database^92^ release 109 with function *DRAM.py annotate* using the default settings. Pathways were evaluated by assessing KOs in KEGG. A list of selected KOs can be found at (**Supplementary Data File 5**), and the DRAM output is available in (**Supplementary Data File 6**). Additionally, pathways, such as the RuMP cycle, were evaluated in MetaCyc^93^. Hydrogenases were annotated by the HydDB webtool^94^. The subunits of dissimilatory sulfate reduction complex (DsrMKJOP) were determined by downloading HMMs from NCBI with accessions NF045798.1 (DsrP), NF045797.1 (DsrO), NF045796.1 (DsrK), NF038037.1 (DsrM), and NF038038.1 (DsrJ). No DsrD could be detected based on generating HMM-profile of a sequence alignment from pfam entry pfam08679. HMM searches were performed using HMMER^95^ v.3.3.2. As the DsrAB can be both oxidative and reductive, we assessed the DrsAB phylogeny. From DRAM, the translated gene sequences annotated as K11180 and K11181 were assessed against DrsAB downloaded from^96^. Sequences were dereplicated at 90% amino acid identity using usearch^75^ v11.0.667 option –cluster_fast, multiple sequence alignment was performed with MAFFT^76^ v7.490, and trimmed at 20% representation with trimAl^77^ v1.5.0. Protein phylogeny was inferred by generating a tree with IQ-TREE^78^ v. 2.1.2, with the ultrafast bootstrap approximation option using 1000 iterations, using translation model WAG. Metabolic potential was parsed and displayed in R^84^ v4.4.0, with packages tidyverse^86^ v2.0.0, readxl^97^ v1.4.3, vroom^98^ v1.6.5, ggplot2^99^ v4.0.0, ggtree^100^ v3.12.0, ape^101^ v5.8.1, ggh4x^102^ v0.2.8 and treeio^103^ v1.28.0. Metabolic potential was summarised in main figures and is displayed in full in (**Supplementary Note 2**).

### Screening of soil samples for oxidation rate experiments

Soils with high taxonomic abundance of selected MAGs were sampled and screened for further analysis. All soils were sampled in depths 0-10 cm and 11-20 cm. Samples were collected in triplicates and mixed. DNA was extracted using the DNeasy PowerSoil Pro kit (QIAGEN, 47016). Qubit dsDNA HS kit (Thermo Fisher, Q33231) with a Qubit 3.0 fluorometer (Thermo Fisher) and NanoDrop One spectrophotometer (Thermo Fisher) was used to determine DNA concentration and quality. Extracted DNA was prepared with the Rapid barcoding SQK-RBK114-96 (Oxford Nanopore). The library was loaded onto a FLO-PRO114M flowcell and sequenced on PromethION PCAMP205 with Super-accurate basecalling v4.3.0, 400 bps. Data was collected with MinKNOW v24.06.15, and basecalled with Dorado v.7.4.14. Reads were filtered to Phred Quality Q10 and minimal read length of 200bp using chopper^104^ v0.9.1 and SeqKit^81^ v2.0.0. The filtered reads were mapped against MFD dereplicated genomes (95% ANI) using CoverM^105^ v0.7.0 with options --min-read-aligned-percent 0.75 --min-read-percent-identity 0.95, --min-covered-fraction 0,-p minimap2-ont.

### Methane oxidation rate experiments

For oxidation rate experiments, soil samples were taken from a Beech forest in Fussingø, Randers (0-10 cm), a Deciduous forest in Aggerholm Søbad, Silkeborg (0-10 cm) and a Xeric sand calcareous grassland from Dokkedal, Mulbjerge (11-20 cm). 20 g of soil was transferred into 250 mL Duran pressure resistant glass bottles. The bottle headspace was flushed for 5 minutes with synthetic air (400 p.p.m.v CO_2_ (HiQ, AGA, Sweden)) using a gas manifold. Before reaching the bottle, the synthetic air was directed through sterile dH_2_O to prevent drying of the soils. The headspace pressure was adjusted to ∼1,08 bar. Finally, 0.5 mL of 1000 ppm methane in nitrogen (HiQ, AGA, Sweden) was added to give a final concentration of approximately 3.5 ppm methane. Samples were incubated at 15°C during the experiment. A negative control was created by autoclaving the soil, and performing the same procedure of flushing, adjusting pressure and adding 0.5 mL of 1000 ppm methane. For each measurement, 2 mL samples of headspace were taken using a gas-tight syringe and injected into a gas chromatograph (Thermo Scientific TRACE 1300, Thermo Fisher Scientific, Waltham, Massachusetts, USA) using the same setup as described in^27^. A 2.5 p.p.m.v CH_4_, 2.5 p.p.m.v H_2_, and 2.5 p.p.m.v CO in N_2_ (HiQ, AGA, Sweden) gas was used to create standard curves. Pressure was measured at the beginning and end of the experiment using a manometer (LEO1, Keller, Winterthur, Switzerland). The amount of methane was determined by applying the ideal gas law. Calculations were adjusted to account for 2 mL gas removal for each sampling point. The rate constants were calculated in R using the integrated rate law for first order reactions. The oxidation rates per gram soil were calculated considering a temperature of 15°C, an absolute pressure of 1.013 bar, a volume of 250 mL, and a relative methane concentration of 1.93 ppm.

### Sequencing of soil post oxidation rate experiments

Following the oxidation rate experiments, soils were frozen and stored at-18°C. DNA was extracted using the DNeasy PowerMax Soil Kit (Qiagen, 12988-10). DNA concentrations were measured with Qubit 3.0 fluorometer (Thermo Fisher Scientific, USA), using dsDNA HS kit (#Q33231, Thermo Fisher Scientific, USA). DNA purity was measured with NanoDrop One Spectrophotometer (Thermo Fisher Scientific, USA). The size of DNA fragments were assessed with Agilent 2200 Tapestation (Agilent Technologies, USA) and Genomic DNA ScreenTapes (#5067-5365, Agilent Technologies, USA). Extracted DNA was prepared for flow cell loading using Ligation Sequencing Kit V14 (SQK-LSK114), and sequenced on PromethION 2 Solo with flowcell FLO-PRO114M. Basecalling was performed with super-accurate basecalling 400bps with Dorado v7.9.8 and MinKNOW v25.05.14. Reads were filtered to be above a Quality Phred score of 10 and read length of 100 bp using chopper^104^ v0.9.1^104^.

Assembly and binning of soil samples was performed using mmlong2 (v1.2.1, https://github.com/Serka-M/mmlong2). Briefly, the long-read metagenomes were assembled with Myloasm^106^ v0.2.0, and automatically curated for misassemblies using rust-anvio-mis (v1.0, https://github.com/bluenote-1577/rust-anvio-mis), which a Rust-based implementation of the Anvio misassembly screening workflow^107^. Contigs that were reported as circular by Myloasm and featured no evidence of misassembly were considered circular and used for cMAG extraction. The remaining contigs were binned with MetaBAT2^108^ v2.18,SemiBin2^109^ v2.2.0, VAMB^110^ v5.0.4 and COMEBin^111^ v1.0.4, using Binette^112^ v1.1.2 to recover the ensemble non-redundant MAGs. Recovered MAGs were dereplicated against the MFD catalogue using dRep^113^ v3.6.2 with standard settings.

## Etymology

### Description of the genus *Methylocalcaria* gen. nov

Methylocalcaria (Me.thy.lo.cal.ca’ri.a. N.L. neut. n. *methȳlum*, methyl group; N.L. masc.. n. *calcarius*, of or pertaining to lime, denoting the presence of this genus in calcareous habitats; N.L. fem. n. *Methylocalcaria*, an organism associated with methyl metabolism in calcareous habitats) The description of this genus is based on three circular high quality MAGs. The type species of the genus is M. *scaenica.* The former genus is ‘JACCXJ01’ in GTDB taxonomy.

Family: Proposed *Candidatus* Methyloligotrophaceae^31^ (former ‘JACCXJ01’ in GTDB taxonomy). Order: Proposed *Candidatus* Methyloligotrophales^31^ (former ‘JACCXJ01’ in GTDB taxonomy).

### Description of the species Methylocalcaria scaenica sp. nov

scaenica (scae’ni.ca. L. fem. adj. *scaenica*, scenic, picturesque, referring to the recovery of the species from a scenic location in calcareous environments). The type genome is represented by genomic assembly GCA_977880055.1, a high quality circular MAG.

### Description of the species Candidatus Methylocalcaria obscura sp. nov

obscura (obs.cu’ra. L. fem. adj. *obscura*, hidden, unseen, referring to its occurrence in only a few restricted habitats). The type genome is represented by genomic assembly GCA_977879555.1, a high quality circular MAG.

### Description of the species Candidatus Methylocalcaria morainei sp. nov

morainei (mo.rai’nei. N.L. gen.. n.. morainei, of or from a moraine). The type genome is represented by genomic assembly GCA_977879905.1, a high quality circular MAG.

### Description of the species Candidatus Methyloligotropha pratensis sp. nov

pratensis (pra.ten’sis. L. fem. adj. pratensis, found in meadows/grassland)

The type genome is represented by genomic assembly GCA_963764725.1, a high quality MAG. Genus: *Candidatus* Methyloligotropha (former ‘JACCXJ01’ in GTDB taxonomy), as proposed in^31^

### Description of the species: Methylocapsa regalis sp. nov

regalis (re.ga’lis. L. fem. adj. regalis, royal, referring to recovery from a forest near Fussingø Castle and high abundance in forest soils). The type genome is represented by genomic assembly GCA_963706215.1, a high quality circular MAG.

### Description of the species Methylocapsa arrakensis sp. nov

arrakensis (ar.ra.ken’sis. N.L. fem. adj. *arrakensis*, of or belonging to Arrakis, the desert planet in Frank Herbert’s Dune series, referring to the abundance of the species in dry habitats). The type genome is represented by genomic assembly GCA_977878845.1, a high quality circular MAG.

### Description of the species Methylocapsa fremenorum sp. nov

fremenorum (fre.men.o.rum. N.L. gen. pl. *fremenorum*, of the Fremen, of or belonging to the Fremen, the desert-dwelling people in Frank Herbert’s *Dune* series, referring to the recovery of the species from a humid dune habitat, inspired by the Fremen dream of conserving and celebrating water. The type genome is represented by genomic assembly GCA_963809115.1, a high quality MAG.

### Description of the species Methylocapsa caladansis sp. nov

caladansis (ca.la.dan’sis. N.L. fem. adj. *caladansis*, of or belonging to Caladan, the lush and wet planet in Frank Herbert’s Dune series, referring to the abundance of the species in humid habitats).The type genome is represented by genomic assembly GCA_963818875.1, a high quality MAG. This is replacing the medium quality genome GCA_003162995.1, *Methylocapsa* sp003162995.

### Description of the species Methylocystis lacustris sp. nov

lacustris (la.cus’tris. N.L. fem. adj. lacustris, from a lake, referring to high abundance in lake sediments). The type genome is represented by the genomic assembly GCA_963717565.1 designated as high quality.

### Description of the species Methylocystis urbana sp. nov

urbana (ur’ba.na. L. fem. adj. urbana, “urban, of a city”, referring to abundance in urban waters on Sjælland). The type genome is represented by the genomic assembly GCA_963706255.1 designated as high quality.

### Description of the species *Rhodomicrobium methylothiovorans* sp. nov

*methylothiovorans* (me.thy.lo.thi.o.vo’rans. N.L. neut. n. *methȳlum*, methyl group; Gr. neut. n. *theîon*, sulfur; L. pres. part. *vorans*, devouring, eating; N.L. neut. part. adj. *methylothiovorans* referring to genomic potential to oxidize methane and reduced sulfur compounds). The type genome is represented by the genomic assembly GCA_963817365.1 designated as high quality.

### Description of the genus Aliimethylobacter gen. nov

*Aliimethylobacter* (A.li.i.me.thy.lo.bac’ter. L. masc. pron. *alius*, another, different; N.L. masc. n. Aliimethylobacter, the other Methylobacter). The type species of the genus is *Aliimethylobacter tundripaludum*.

### Description of the species *Aliimethylobacter* imbricus

imbricus (im’bri.cus. L. masc. adj. *imbricus*, pertaining to rain, rainy, referring to occurrence in urban enclosed waters and rainwater basins). The type genome is represented by the genomic assembly GCA_963793545.1 designated as high quality.

Genus: *Aliimethylobacter* (former ‘Methylobacter_A’ in GTDB taxonomy).

Genomes will be submitted for registration at SeqCode (<u>SeqCode Registry</u>). The complete protologue table is provided in (**Supplementary Data File 3**).

