## Supplementary Material for "A national baseline for methane sink habitats and methanotroph diversity"

*Corresponding Author

**Correspondence**

Caitlin M. Singleton

Center for Microbial Communities, Department of Chemistry and Bioscience,

Aalborg University, Fredrik Bajers Vej 7H, 9220 Aalborg, Denmark

**Table of content**


Extended Data Figures 1-13

Supplementary Data Files 1-6

Supplementary Notes 1 & 2

Supplementary References

**Extended data figures:**

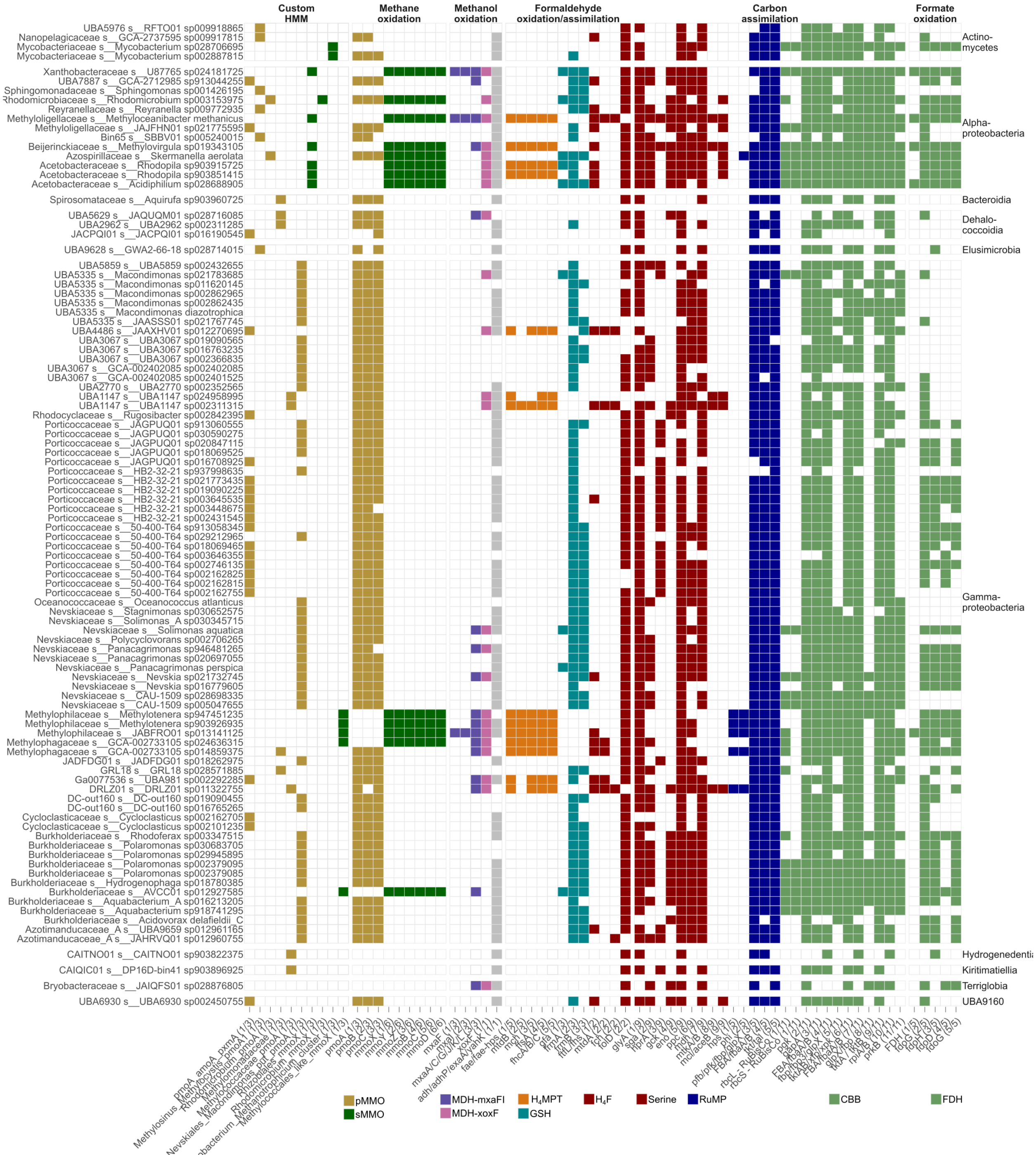


**Extended Data Figure 1: Genomic potential of putative methanotrophs.** Methane related metabolic potential of GTDB r220 genomes with putative *pmoA* or *mmoX* sequences quantified with our GraftM packages (Custom HMM) and DRAM. Genome taxonomy is indicated by family and species, and faceted by class.

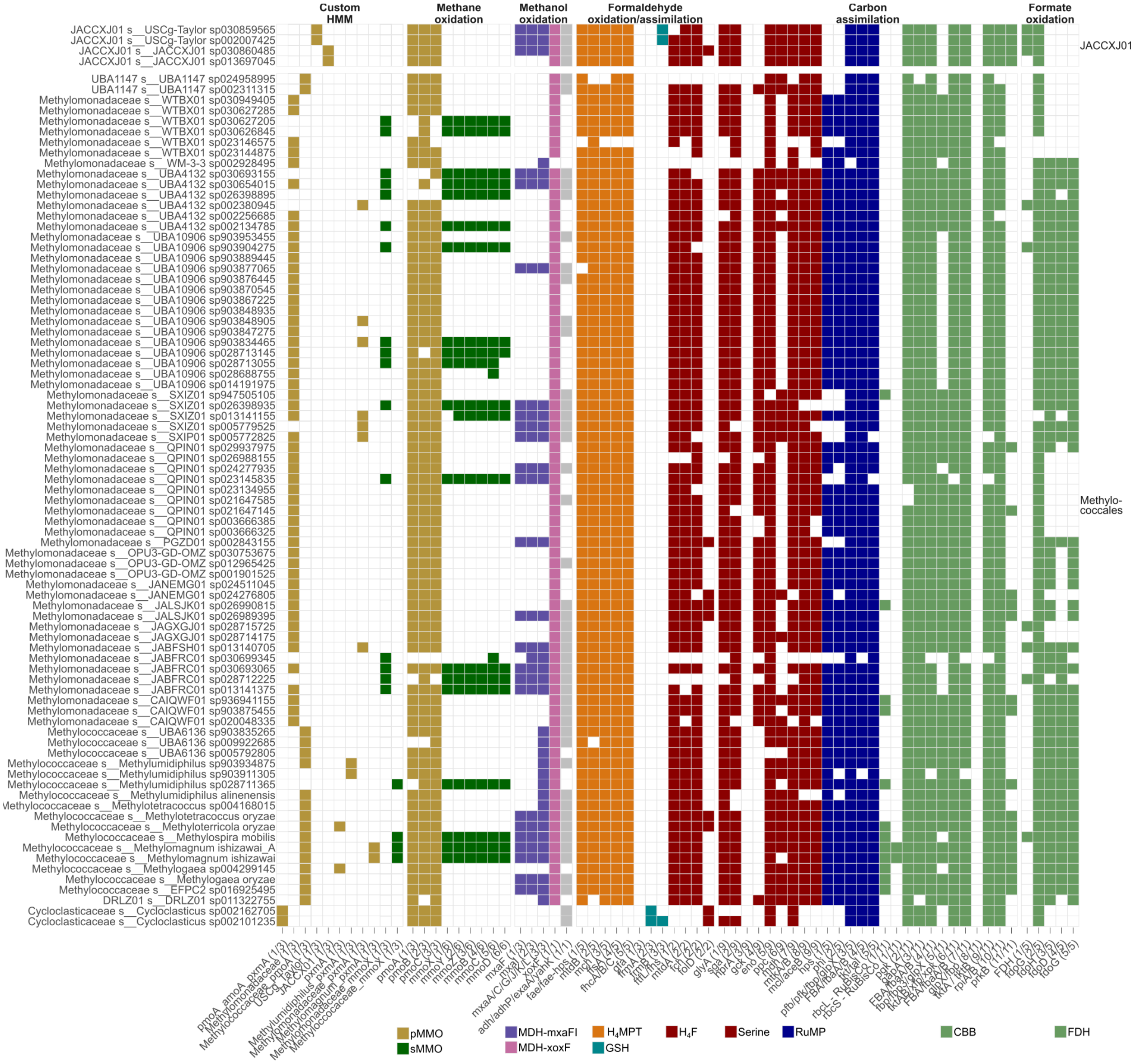


**Extended Data Figure 2: Genomic potential of putative Methylococcales methanotrophs.** Methane related metabolic potential of Methylococcales and JACCXJ01 GTDB r220 genomes with putative *pmoA* or *mmoX* sequences quantified with our GraftM packages (Custom HMM) and DRAM. Genome taxonomy is indicated by family and species, and faceted by class.

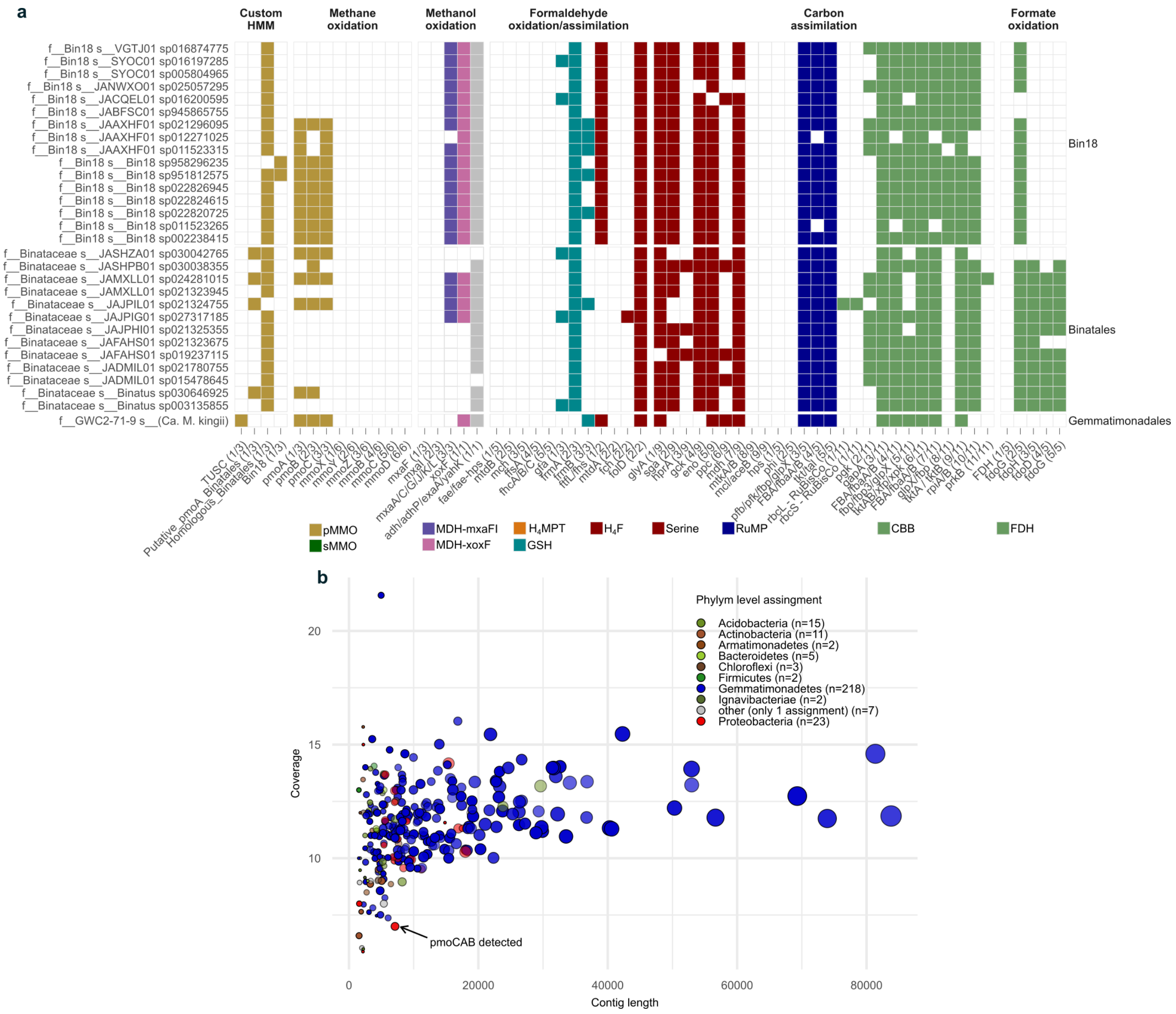


**Extended Data Figure 3: Genomic potential and contamination of putative Binatia and TUSC genomes. a)** Methane related metabolic potential of BinatiaGTDB r220 genomes and *Ca.* M. kingii with putative *pmoA* sequences quantified with our GraftM packages (Custom HMM) and DRAM. Genome taxonomy is indicated by family and species, and faceted by class. **b)** Taxonomy, coverage and length of contigs in *Ca.* M. kingii metagenome assembled genome. The contigs are scaled in size by number of coding sequences, and are coloured by the phylum level assignment from GUNC v1.0.6[1]. The coverage and length was obtained from the fasta file of “DMC_bin26”.

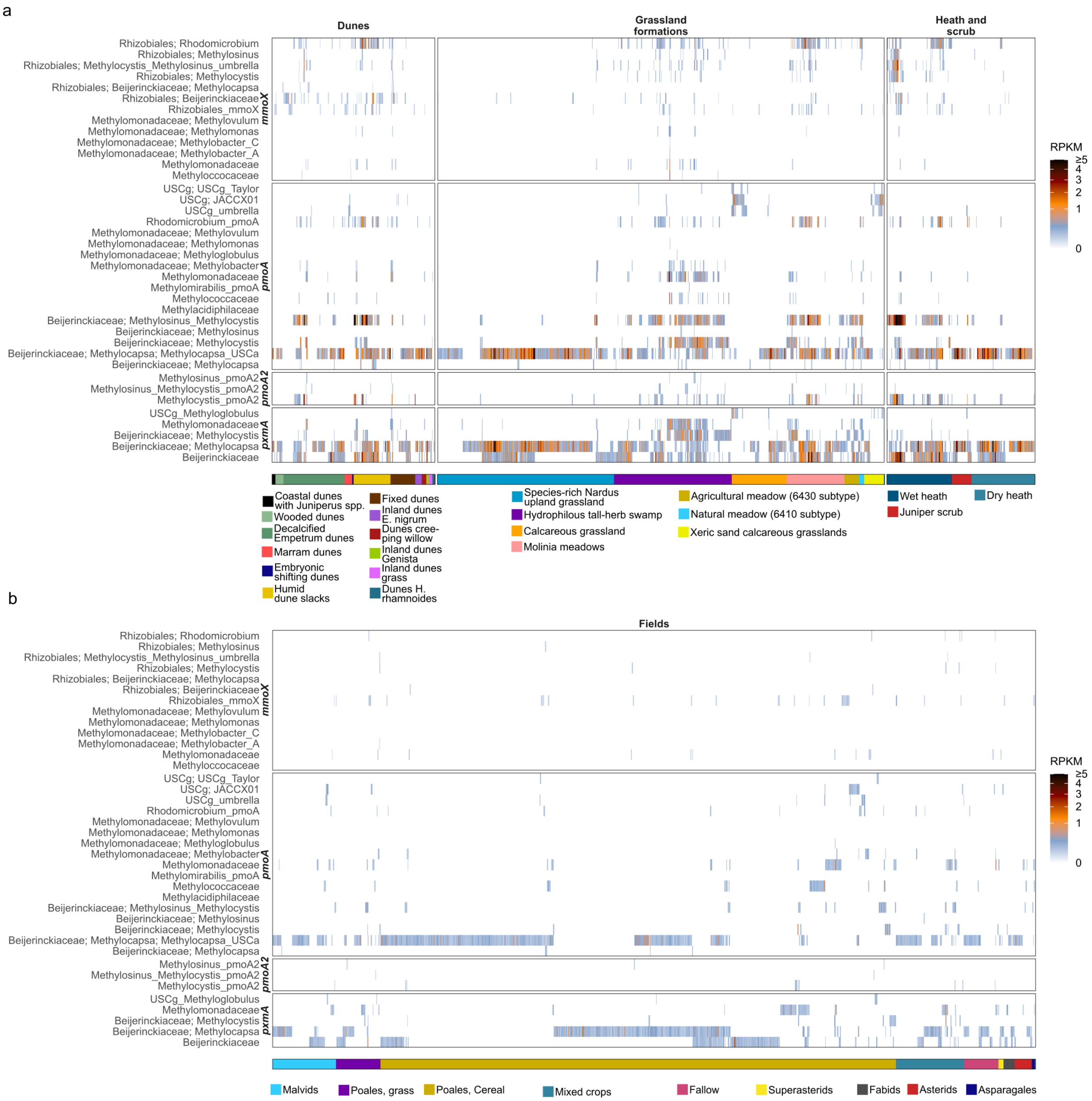


**Extended Data Figure 4: Relative gene abundance of *pmoA* and *mmoX* in selected habitats:** **a)** Distribution of methanotrophs marker genes identified with our *pmoA* and *mmoX* GraftM packages in dunes, grasslands and heaths, displayed on the MFD ontology level 3 (MFDO3). Samples are coloured by the number of reads (Reads Per Kilobase Million (RPKM)) assigned to each gene-phylogenetic group. Samples are clustered with hierarchical clusters within each MFDO3 habitat. The bottom colour panel indicates the MFDO3 habitat. **b)** Distribution of methanotrophs marker genes in agricultural samples, clustered within each crop type (MFDO2).

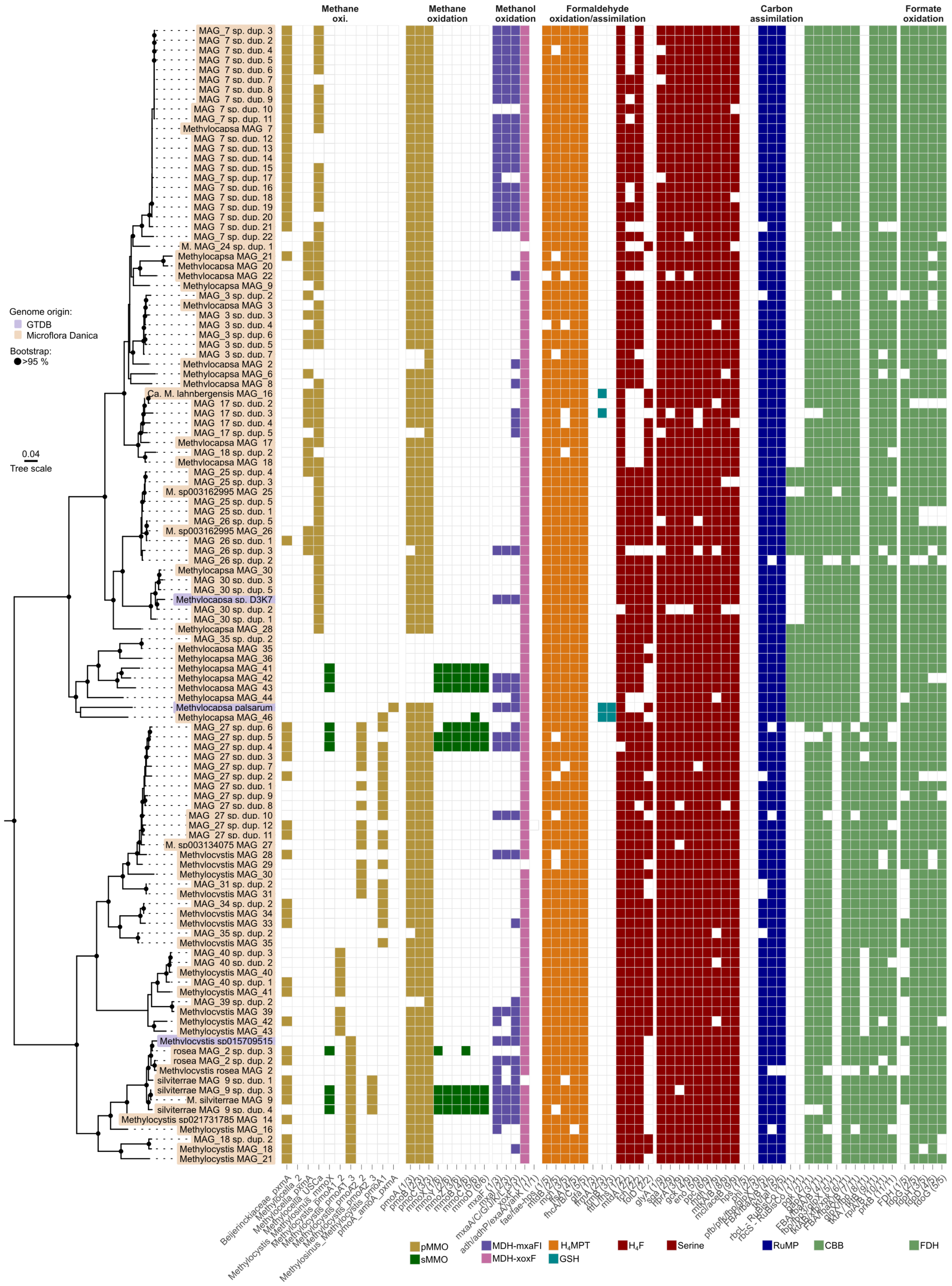
Methane oxi.
Methane oxidation
Methanol oxidation
Formaldehyde oxidation/assimilation
Carbon assimilation
Formate oxidation
Genome origin:
GTDB
Microflora Danica
Bootstrap:
>95 %
0.04
Tree scale
MAG_7 sp. dup. 3
MAG_7 sp. dup. 2
MAG_7 sp. dup. 4
MAG_7 sp. dup. 5
MAG_7 sp. dup. 6
MAG_7 sp. dup. 7
MAG_7 sp. dup. 8
MAG_7 sp. dup. 9
MAG_7 sp. dup. 10
MAG_7 sp. dup. 11
Methylocapsa MAG_7
MAG_7 sp. dup. 12
MAG_7 sp. dup. 13
MAG_7 sp. dup. 14
MAG_7 sp. dup. 15
MAG_7 sp. dup. 17
MAG_7 sp. dup. 16
MAG_7 sp. dup. 18
MAG_7 sp. dup. 19
MAG_7 sp. dup. 20
MAG_7 sp. dup. 21
MAG_7 sp. dup. 22
M. MAG_24 sp. dup. 1
Methylocapsa MAG_21
Methylocapsa MAG_20
Methylocapsa MAG_22
Methylocapsa MAG_9
MAG_3 sp. dup. 2
Methylocapsa MAG_3
MAG_3 sp. dup. 3
MAG_3 sp. dup. 4
MAG_3 sp. dup. 6
MAG_3 sp. dup. 5
MAG_3 sp. dup. 7
Methylocapsa MAG_2
Methylocapsa MAG_6
Methylocapsa MAG_8
Ca. M. lahnbergensis MAG_16
MAG_17 sp. dup. 2
MAG_17 sp. dup. 3
MAG_17 sp. dup. 4
MAG_17 sp. dup. 5
Methylocapsa MAG_17
MAG_18 sp. dup. 2
Methylocapsa MAG_18
MAG_25 sp. dup. 4
MAG_25 sp. dup. 3
M. sp003162995 MAG_25
MAG_25 sp. dup. 5
MAG_25 sp. dup. 1
MAG_26 sp. dup. 5
M. sp003162995 MAG_26
MAG_26 sp. dup. 1
MAG_26 sp. dup. 3
MAG_26 sp. dup. 2
Methylocapsa MAG_30
MAG_30 sp. dup. 3
MAG_30 sp. dup. 5
Methylocapsa sp. D3K7
MAG_30 sp. dup. 2
MAG_30 sp. dup. 1
Methylocapsa MAG_28
MAG_35 sp. dup. 2
Methylocapsa MAG_35
Methylocapsa MAG_36
Methylocapsa MAG_41
Methylocapsa MAG_42
Methylocapsa MAG_43
Methylocapsa MAG_44
Methylocapsa palsarum
Methylocapsa MAG_46
MAG_27 sp. dup. 6
MAG_27 sp. dup. 5
MAG_27 sp. dup. 4
MAG_27 sp. dup. 3
MAG_27 sp. dup. 7
MAG_27 sp. dup. 2
MAG_27 sp. dup. 1
MAG_27 sp. dup. 9
MAG_27 sp. dup. 8
MAG_27 sp. dup. 10
MAG_27 sp. dup. 12
MAG_27 sp. dup. 11
M. sp003134075 MAG_27
Methylocystis MAG_28
Methylocystis MAG_29
Methylocystis MAG_30
MAG_31 sp. dup. 2
Methylocystis MAG_31
MAG_34 sp. dup. 2
Methylocystis MAG_34
Methylocystis MAG_33
MAG_35 sp. dup. 2
Methylocystis MAG_35
MAG_40 sp. dup. 3
MAG_40 sp. dup. 2
Methylocystis MAG_40
MAG_40 sp. dup. 1
Methylocystis MAG_41
MAG_39 sp. dup. 2
Methylocystis MAG_39
Methylocystis MAG_42
Methylocystis MAG_43
Methylocystis sp015709515
rosea MAG_2 sp. dup. 3
rosea MAG_2 sp. dup. 2
Methylocystis rosea MAG_2
silviterrae MAG_9 sp. dup. 1
silviterrae MAG_9 sp. dup. 3
M. silviterrae MAG_9
silviterrae MAG_9 sp. dup. 4
Methylocystis sp021731785 MAG_14
Methylocystis MAG_16
MAG_18 sp. dup. 2
Methylocystis MAG_18
Methylocystis MAG_21
pMMO
sMMO
MDH-mxaFI
MDH-xoxF
H4MPT
GSH
H4F
Serine
RuMP
CBB
FDH

**Extended Data Figure 5: Metabolic potential and genome phylogeny of *Methylocapsa* and *Methylocystis* MAGs.** Methane related metabolic potential of *Methylocapsa* and *Methylocystis* with putative *pmoA* or *mmoX* sequences quantified with our GraftM packages (Custom HMM) and DRAM. The phylogenomic tree of *Methylocapsa* and *Methylocystis* spp. from our recovered MAGs and selected genomes from GTDB is based on genomes filtered at >80% completeness and <10% contamination.

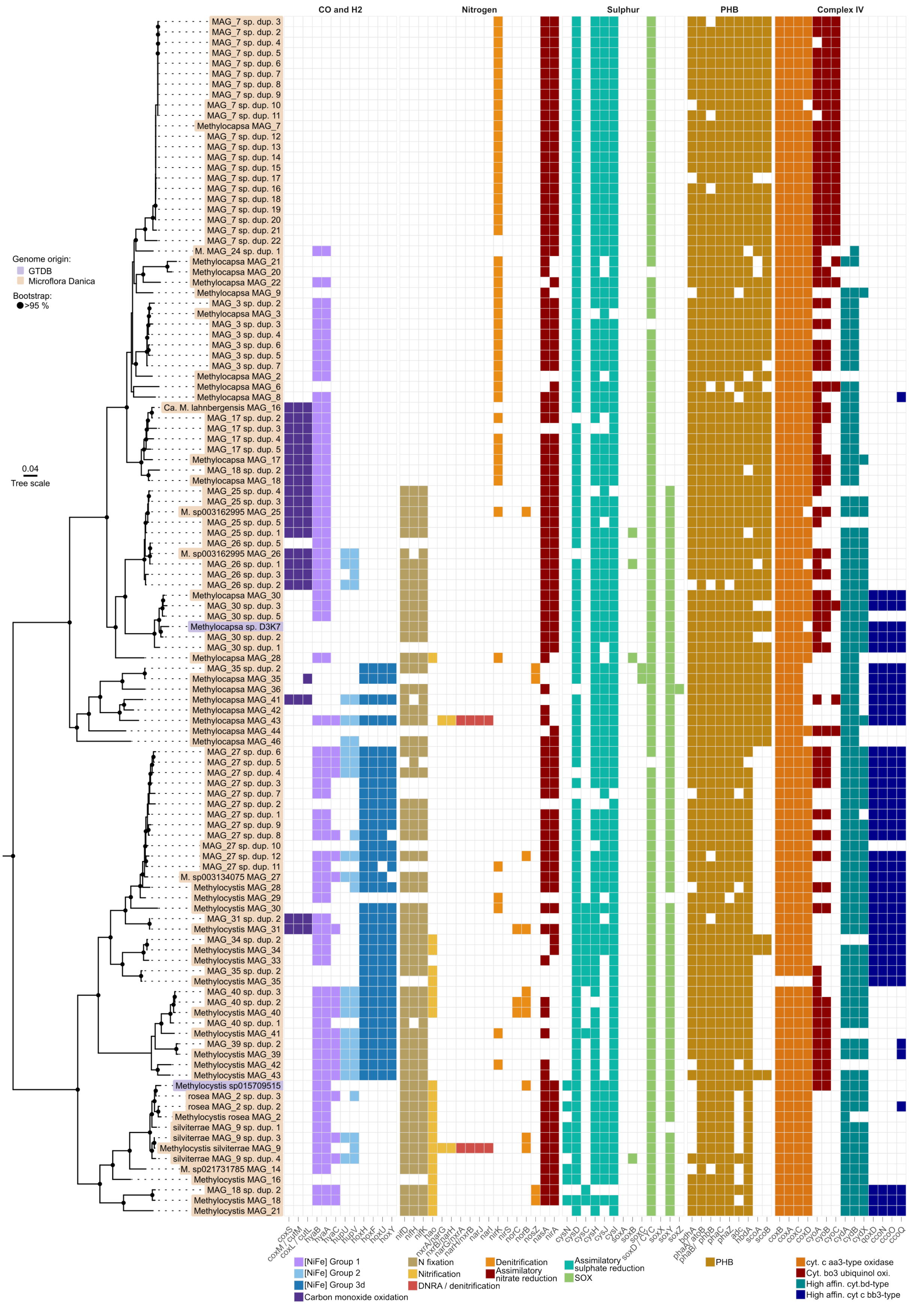
CO and H2
Nitrogen
Sulphur
PHB
Complex IV
Genome origin:
GTDB
Microflora Danica
Bootstrap:
>95 %
0.04
Tree scale
[NiFe] Group 1
[NiFe] Group 2
[NiFe] Group 3d
Carbon monoxide oxidation
N fixation
Nitrification
DNRA / denitrification
Denitrification
Assimilatory nitrate reduction
Assimilatory sulphate reduction
SOX
PHB
cyt. c aa3-type oxidase
Cyt. bo3 ubiquinol oxi.
High affin. cyt.bd-type
High affin. cyt c bb3-type

**Extended Data Figure 6: Metabolic potential and alternative metabolism of *Methylocapsa* and *Methylocystis* MAGs.** Phylogenomic tree of *Methylocapsa* and *Methylocystis* spp. from our recovered MAGs and selected genomes from GTDB. All genomes have been filtered at >80% completeness and <10% contamination. Selected metabolic modules assigned with DRAM and based on KOs are displayed for oxidation of carbon monoxide, oxidation of hydrogen, along with nitrogen and sulphur metabolisms. Additionally, genes associated with PHB storage are shown, along with the complex IV for transfer of electrons to the terminal electron acceptor.

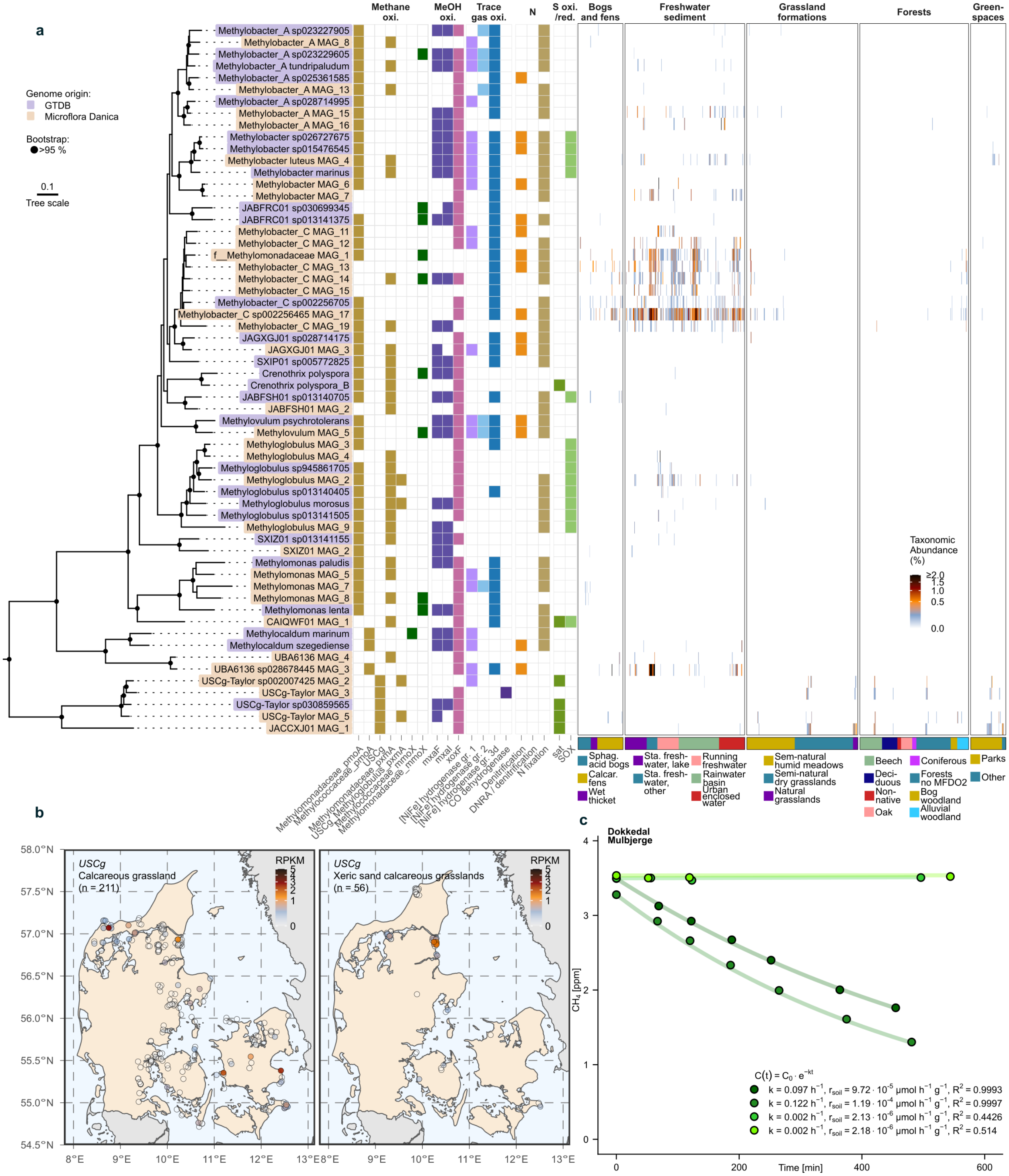


**Extended Data Figure 7: Quantification of and characterisation of gammaproteobacterial known and putative methanotrophs. a.** Phylogenomic tree of selected known and putative methanotrophs within Gammaproteobacteria from GTDB and MFD recovered MAGs. Genomes have been dereplicated at 95% ANI. Bootstraps with above 95% support (UFB, 1000 replications) are indicated by a black circle. Associated with each genome is the phylogeny of *pmoA, pxmA*

and *mmoX,* along with metabolic potential for methanol oxidation, trace gas oxidation ($H_2$ and CO), nitrogen fixation, denitrification, and sat and SOX complexes in sulphur oxidation. Additionally, each genome has been quantified (sylph), and the taxonomic abundance is shown across the major natural habitats. Habitats are clustered (hclust) within MFDO2. **b.** Quantification of USCγ gene abundance (RPKM) in Calcareous grassland and Xeric sand calcareous grasslands samples in Denmark (map from EuroGeographics). **c.** Oxidation rate experiment of a selected soil from Dokkedal, Mulbjerge (Xeric sand calcareous grasslands). Headspace concentrations of methane were measured from four replicates of the soil. A linear model (function lm()) of the natural log of methane concentrations was calculated to find first order decay (k, $h^{-1}$) constant and correlation coefficient.

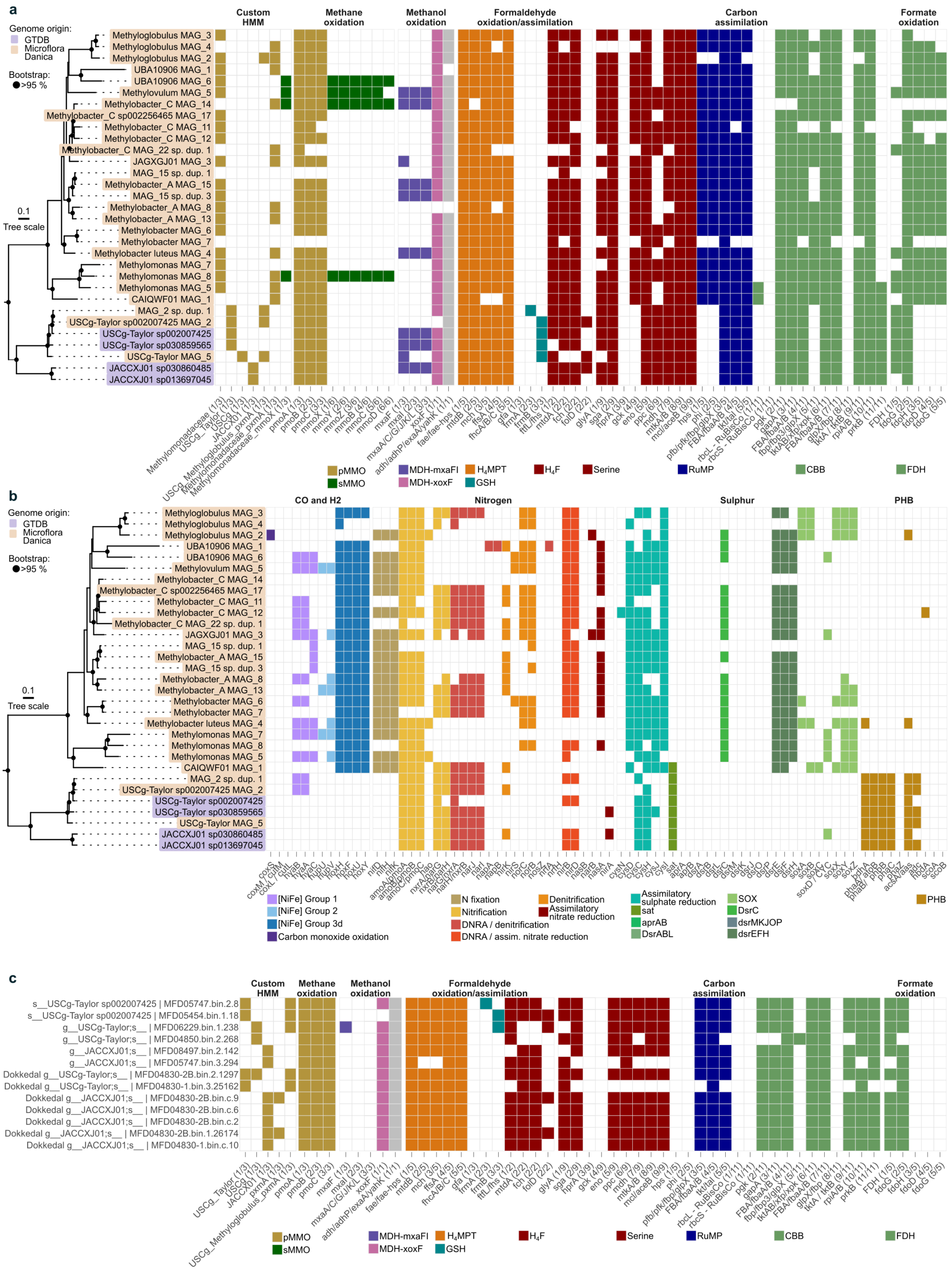


**Extended Data Figure 8: Metabolic potential of gammaproteobacterial methanotrophs.**

**a)** Methane related metabolic potential of gammaproteobacteria with putative *pmoA* or *mmoX* sequences quantified with our GraftM packages (Custom HMM) and DRAM. The phylogenomic tree is based on our recovered MAGs and selected genomes from GTDB filtered at >80% completeness and <10% contamination. **b)** Metabolic potential of gammaproteobacteria from our recovered MAGs and selected genomes from GTDB filtered at >80% completeness and <10% contamination. Shown is metabolic potential related to trace gas oxidation ($H_2$ and CO), nitrogen and sulphur metabolism, and potential to store PHB. **c)** Methane centric metabolic potential of all putative USCγ methanotrophs, with no completeness or contamination filter. The plot includes MAGs recovered from the Dokkedal oxidation rate experiment, denoted by “Dokkedal” in the genome name.

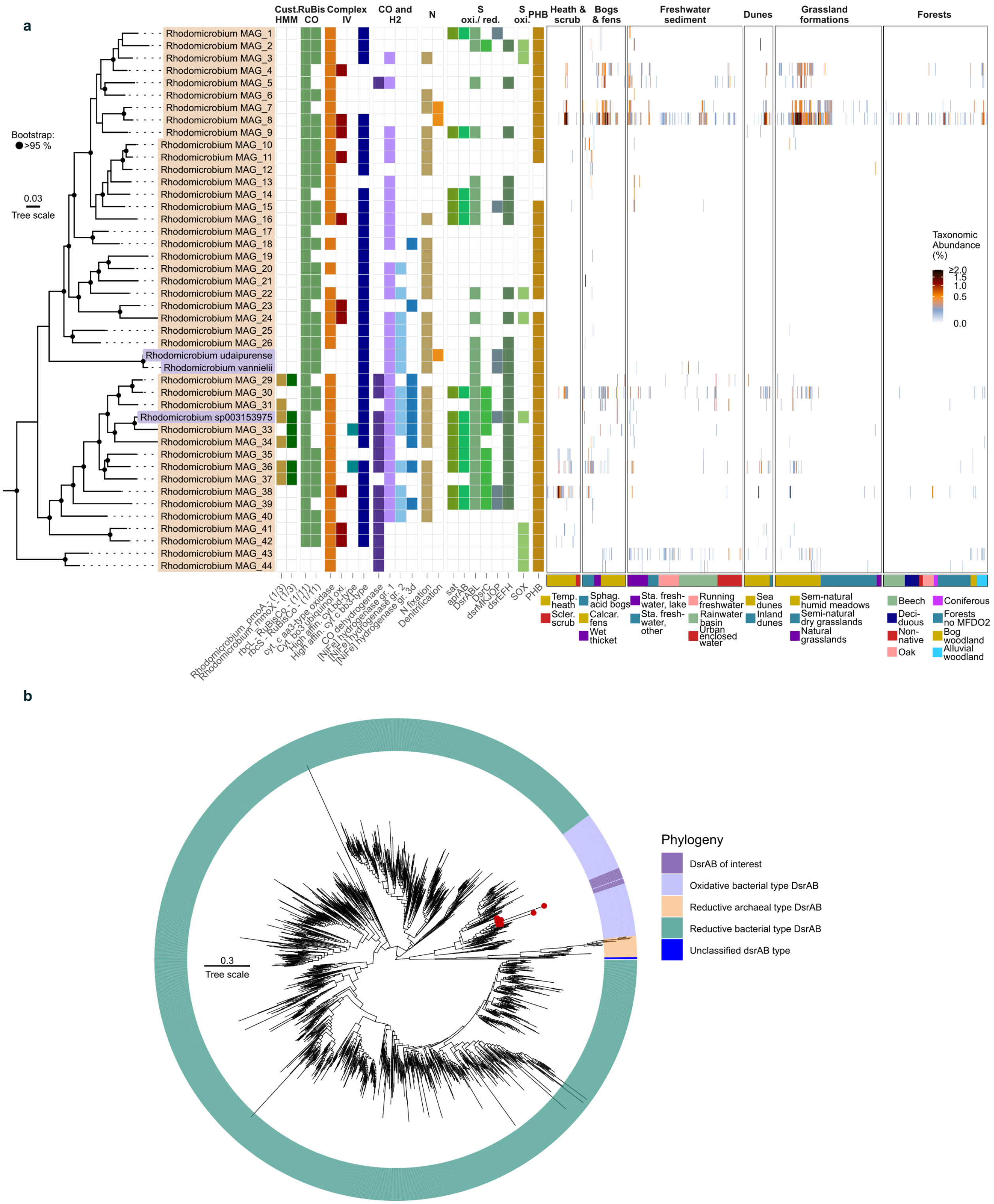


**Extended Data Figure 9: Quantification of and characterisation of *Rhodomicrobium* MAGs.**

**a.** Phylogenomic tree of *Rhodomicrobium* genomes from GTDB and MFD recovered MAGs. Genomes have been dereplicated at 95% ANI. Bootstraps with above 95% support (UFB, 1000 replications) are indicated by a black circle. Associated with each genome is the phylogeny of *pmoA* and *mmoX,* along with metabolic potential for $CO_2$-fixation with RuBisCO, terminal oxidase of the electron transport chain, oxidation of $H_2$ and CO, oxidation and reduction of sulphur compounds, and production of PHB. Additionally, each genome has been quantified (sylph), and the taxonomic abundance is shown across the major natural habitats. Habitats are clustered (hclust) within MFDO2. **b.** Phylogenetic tree of DsrAB. Recovered *dsrA* and *dsrB* from our MAGs were concatenated and phylogeny was inferred based on sequences from [2]. The DsrAB from our MAGs are indicated in red dots and highlighted as part of the oxidative type in dark purple.

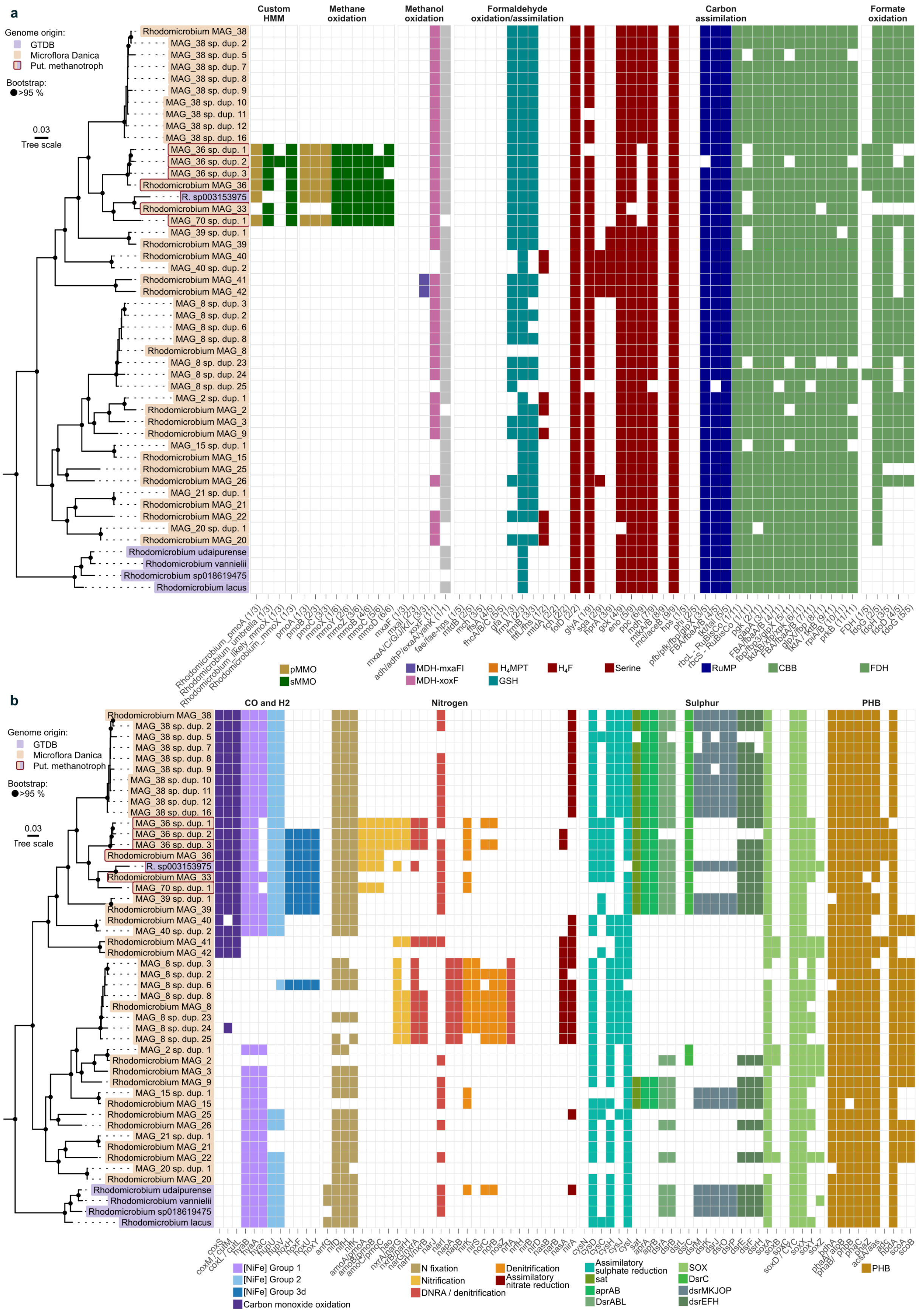
a
Genome origin:
GTDB
Microflora Danica
Put. methanotroph
Bootstrap:
>95 %
0.03
Tree scale
Custom HMM
Methane oxidation
Methanol oxidation
Formaldehyde oxidation/assimilation
Carbon assimilation
Formate oxidation
Rhodomicrobium MAG_38
MAG_38 sp. dup. 2
MAG_38 sp. dup. 5
MAG_38 sp. dup. 7
MAG_38 sp. dup. 8
MAG_38 sp. dup. 9
MAG_38 sp. dup. 10
MAG_38 sp. dup. 11
MAG_38 sp. dup. 12
MAG_38 sp. dup. 16
MAG_36 sp. dup. 1
MAG_36 sp. dup. 2
MAG_36 sp. dup. 3
Rhodomicrobium MAG_36
R. sp003153975
Rhodomicrobium MAG_33
MAG_70 sp. dup. 1
MAG_39 sp. dup. 1
Rhodomicrobium MAG_39
Rhodomicrobium MAG_40
MAG_40 sp. dup. 2
Rhodomicrobium MAG_41
Rhodomicrobium MAG_42
MAG_8 sp. dup. 3
MAG_8 sp. dup. 2
MAG_8 sp. dup. 6
MAG_8 sp. dup. 8
Rhodomicrobium MAG_8
MAG_8 sp. dup. 23
MAG_8 sp. dup. 24
MAG_8 sp. dup. 25
MAG_2 sp. dup. 1
Rhodomicrobium MAG_2
Rhodomicrobium MAG_3
Rhodomicrobium MAG_9
MAG_15 sp. dup. 1
Rhodomicrobium MAG_15
Rhodomicrobium MAG_25
Rhodomicrobium MAG_26
MAG_21 sp. dup. 1
Rhodomicrobium MAG_21
Rhodomicrobium MAG_22
MAG_20 sp. dup. 1
Rhodomicrobium MAG_20
Rhodomicrobium udaipurense
Rhodomicrobium vannielii
Rhodomicrobium sp018619475
Rhodomicrobium lacus
pMMO
sMMO
MDH-mxaFI
MDH-xoxF
H4MPT
GSH
H4F
Serine
RuMP
CBB
FDH
b
CO and H2
Nitrogen
Sulphur
PHB
[NiFe] Group 1
[NiFe] Group 2
[NiFe] Group 3d
Carbon monoxide oxidation
N fixation
Nitrification
DNRA / denitrification
Denitrification
Assimilatory nitrate reduction
Assimilatory sulphate reduction
sat
aprAB
DsrABL
SOX
DsrC
dsrMKJOP
dsrEFH
PHB

**Extended Data Figure 10: Metabolic potential of Rhodomicrobium MAGs. a)** Methane related metabolic potential of *Rhodomicrobium* genomes with putative *pmoA* or *mmoX* sequences quantified with our GraftM packages (Custom HMM) and DRAM. The phylogenomic tree is based on our recovered MAGs and selected genomes from GTDB filtered at >80% completeness and <10% contamination. Putative methanotrophs are indicated with a red outline. **b)** Metabolic potential of *Rhodomicrobium* genomes from our recovered MAGs and selected genomes from GTDB filtered at >80% completeness and <10% contamination. Shown is metabolic potential related to trace gas oxidation ($H_2$ and CO), nitrogen and sulphur metabolism, and potential to store PHB. In both **a** and **b, the** species duplicates of *Rhodomicrobium* MAG_8 (sp. dup. 1, 4, 5, 7, 9, 15, 17, 19, 20, 22, 26, 27, 28) and *Rhodomicrobium* MAG_38 (sp. dup. 3, 4, 6, 8,10, 13, 14, 15) along with genomes *Rhodomicrobium* MAG_10, *Rhodomicrobium* MAG_11, *Rhodomicrobium* MAG_12, *Rhodomicrobium* MAG_13, *Rhodomicrobium* MAG_14, *Rhodomicrobium* MAG_15, *Rhodomicrobium* MAG_16, *Rhodomicrobium* MAG_17, and *Rhodomicrobium* MAG_18 were removed from the plot for readability.

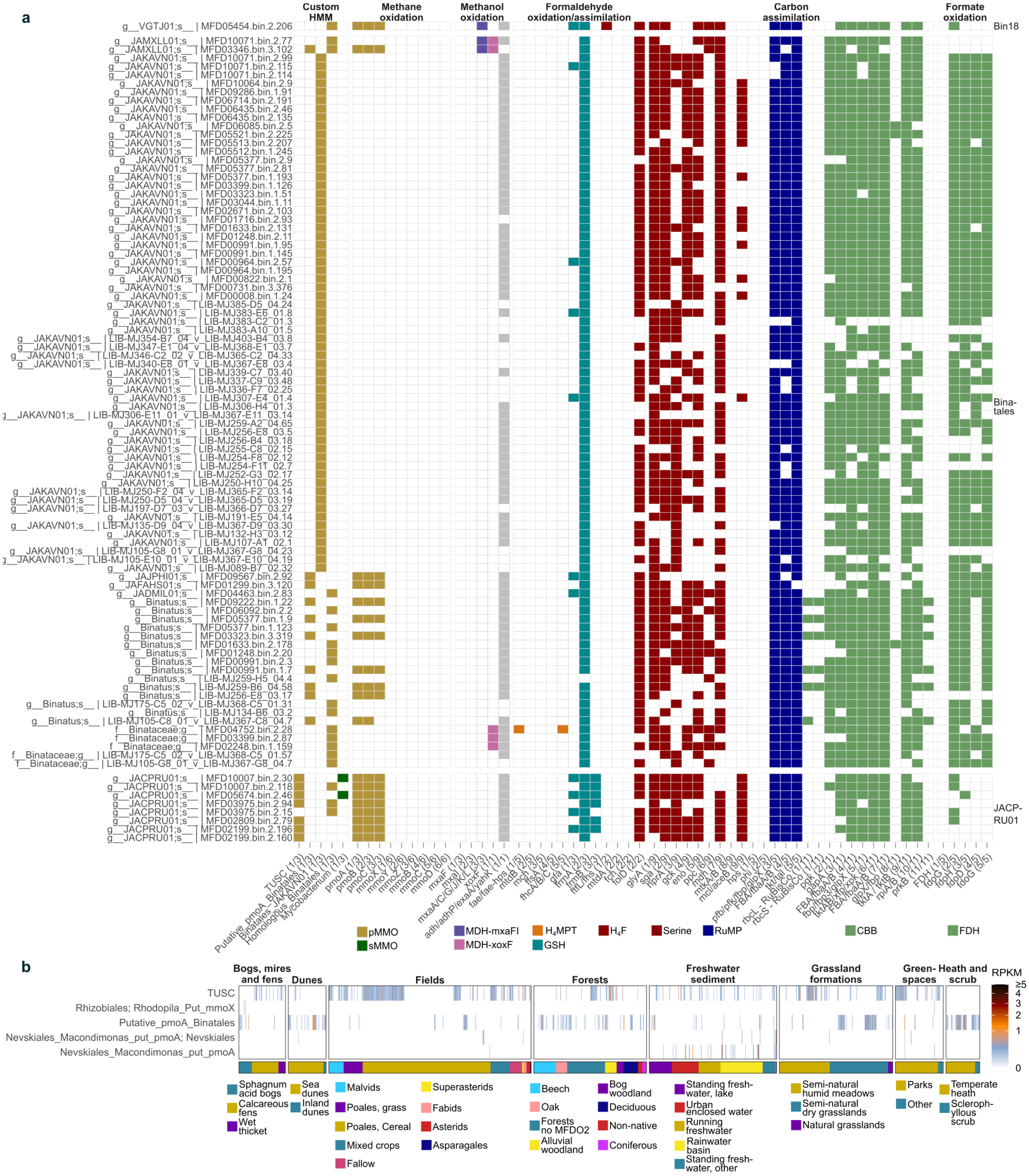


**Extended Data Figure 11: Quantification and metabolic potential of TUSC and Binatales. a)** Methane centric metabolic potential of all Binatia genomes, with no completeness or contamination filter. Genomes are faceted by order, and indicated by specific ID, and lowest possible identification with gtdb-tk (genus or family). **b)** Heatmap of putative *pmoA*-like clades quantified with our GraftM package. Samples are coloured by the number of reads (Reads Per

Kilobase Million (RPKM)) assigned to each gene-phylogenetic group. Samples are clustered with hierarchical clusters within each MFDO2 habitat. The bottom colour panel indicates the MFDO2 habitat.

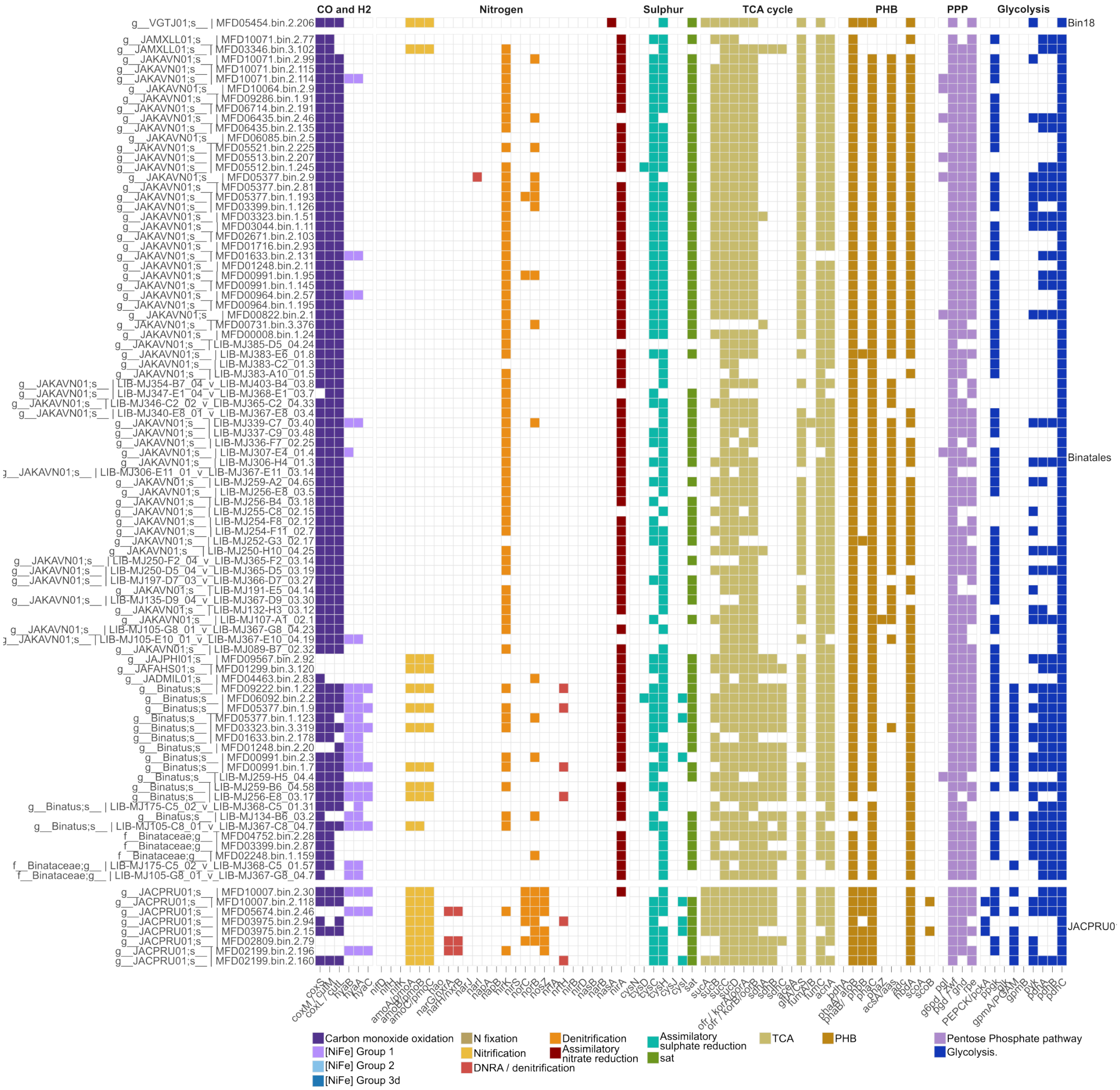


**Extended Data Figure 12: Alternative metabolism of TUSC and Binatales MAGs.** Metabolic potential of Binatia genomes from our recovered MAGs. Shown is metabolic potential related to trace gas oxidation ($H_2$ and CO), nitrogen and sulphur metabolism, tricarboxylic acid cycle (TCA) cycle, potential to store PHB, Pentose Phosphate Pathway (PPP), and glycolysis.

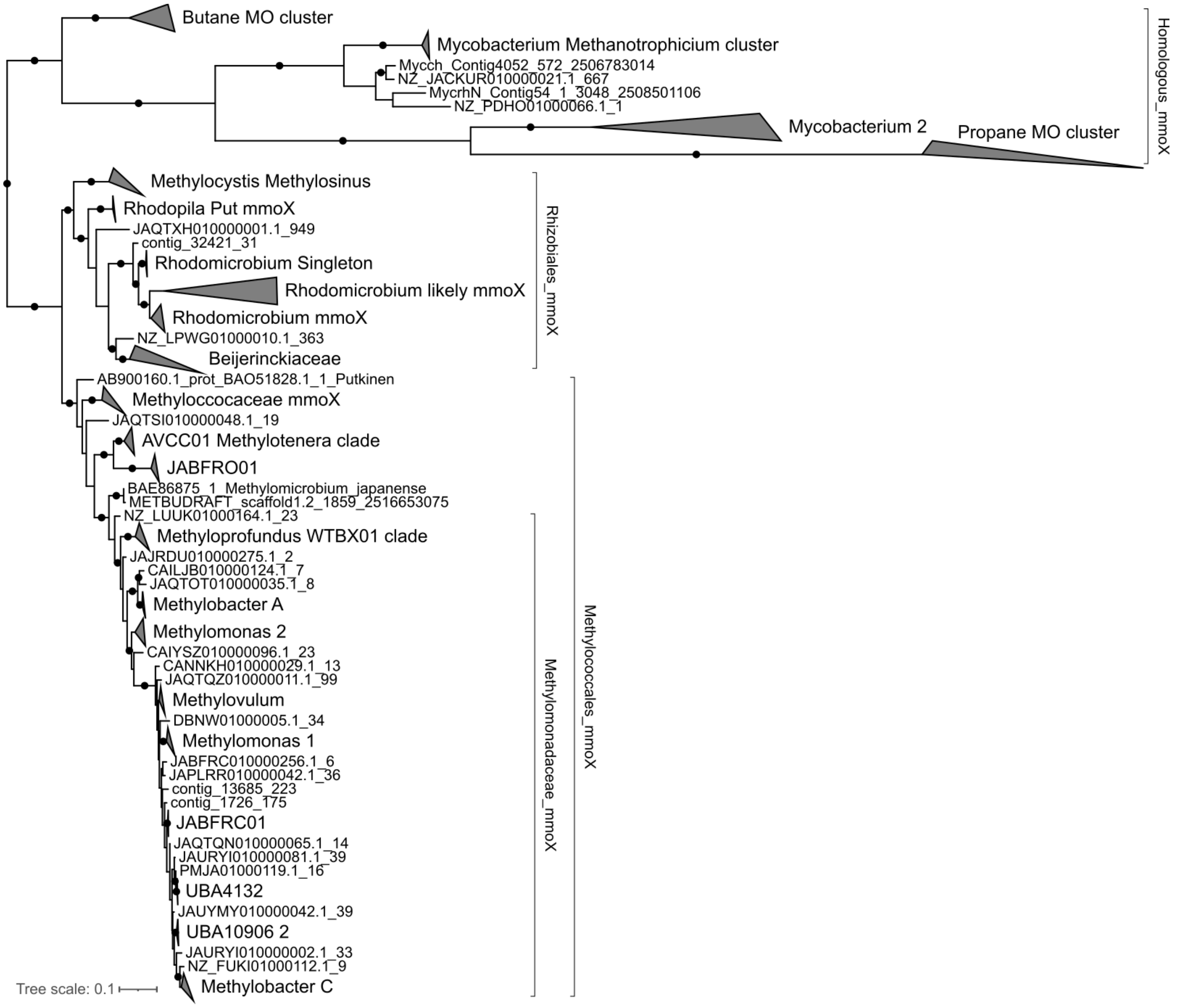


**Extended Data Figure 13: *mmoX* phylogeny.** Protein phylogenetic tree of MmoX sequences used for creation of our custom GraftM package. Clades are depicted as shown in heatmaps and correspond to the GraftM package. Bootstraps with above 95% support (UFB, 1000 replications) are indicated by a black circle.

#### Supplementary Data Files:

**Supplementary Data File 1:** Oxi_rate_mmlong2_bins.tsv. MAG statistics for oxidation rate experiments.

**Supplementary Data File 2:** methanotroph_IDs.tsv. Microflora Danica MAGs analysed here for methanotrophy.

**Supplementary Data File 3:** Protologue_table_methanotrophs.xlsx. Protologue table for proposed names for selected microbial lineages.

**Supplementary Data File 4:** GraftM_output.zip. Output of *pmoA* and *mmoX* GraftM packages.

**Supplementary Data File 5:** KO_KSK_methanotroph_paper.xlsx. A list of KEGG Orthology (KO) identifiers used to perform functional annotation.

**Supplementary Data File 6:** Metabolism_DRAM.7z. Functional annotation of methanotrophs. DRAM functional annotation output for methanotrophs described in this study.

All supplementary data files can be found at GitHub: (https://github.com/KalinkaKnudsen/MFD_methanotrophs_DK/tree/main/data)

### Supplementary Notes

#### Supplementary Note 1: Screening of GTDB for potential methanotrophs

We screened species representatives of GTDB r220 for methanotrophy marker genes using the *pmoA* and *mmoX* graftM packages from Singleton 2018[3], with additional sequences from recent publications. We first evaluated the gene-phylogeny of the potential hits. Next, we assessed the metabolic potential of genomes encoding *pmoA*-like sequences using DRAM[4].

In cases where *pmoA* and *mmoX* sequences were identified in yet-undescribed methanotrophic groups, signs of contamination and genomic potential for methanotrophy were investigated. This included TUSC-*pmoA* sequences in the Gemmatimonadota phylum[5], *Methylococcaceae-pmoA* sequences in phyla Bacteroidota (GCA_903960725.1), Hydrogenedentota (GCA_903822375.1), CSSED10-310 (GCA_016932295.1), o__GRL18 (GCA_028571885.1) and Chloroflexota (f__UBA2962 GCA_002311285.1, GCA_028716085.1), along with *Methylocystaceae-pmoA* sequences in the Actinomycetota phylum (f__Nanopelagicaceae GCA_009917815.1, f__UBA5976 GCA_009918865.1) and alphaproteobacterial orders Reyranellales (GCA_009772935.1), Bin65 (GCA_005240015.1), and Sphingomonadales (GCF_001426195.1), and c__Elusimicrobia;o__UBA1565 (GCA_028714015.1). The *pmoA* gene was found on extremely small contigs (1200-7000 bp) in all above mentioned genomes. The CDS in the given contigs had best hits (blastp against nr) to *Methylococcaceae* or *Methylocystaceae* proteins, while CDS from the surrounding contigs aligned to proteins from other orders or phyla. Metabolic annotation with DRAM revealed missing key enzymes for downstream metabolisation of methane. On this basis, presence of *pmoA* in abovementioned genomes was evaluated to be contamination.

We identified putative pMMOs within the gammaproteobacterial families DRLZ01 and UBA1147, both part of the *Methylococcales* order (**Table 1**). In GTDB r220, only three species were present in the family UBA1147 of which two contained *Methylococcaceae*-like *pmoAs* (GCA_002311315.1 (recovered from a hydrothermal vent) and GCA_024958995.1 (recovered from a Gulf of Mexico water column), while the third species encoded *pmoBC* on the edge of a contig. Two species were classified to the DRLZ01 family, of which one contained a *Methylococcaceae*-like *pmoA* (GCA_011322755.1), while the other (GCA_022600935.1) did not contain any *pmo* genes, and incomplete assimilation and methanol processing pathways. Both families contained no isolates or HQ genomes, and all MAGs encoded partial potential for downstream methanol processing and assimilation, with GCA_011322755 (DRLZ01) encoding all genes for the RuMP cycle, and UBA1147 having only incomplete potential for carbon assimilation (**Extended Data Figure 1**). Within Alphaproteobacteria, *Skermanella aerolata* contained *pmoCAB* operon, *xoxF*-type methanol dehydrogenase, formaldehyde and formate oxidation, and complete CBB cycle (**Extended Data Figure 1**), but methane oxidation activity remains to be determined[6]**.**

Screening GTDB r220 also identified a potential sMMO containing methanotroph within the alphaproteobacterial family *Xanthobacteraceae*, g__U87765 (GCF_024181725.1, recovered from rice roots). The genome is of MIMAG HQ and consists of a single contig, and encodes the entire *mmoXYZBD* operon, calcium-dependent and *xoxF*-type methanol dehydrogenase, formaldehyde oxidation through GSH pathway, along with formate oxidation and assimilation of carbon through CBB cycle (**Extended Data Figure 1**). This is the first reported instance of methanotrophy within the *Xanthobacteraceae* family.

Furthermore, three members of the *Acetobacteraceae* family (Alphaproteobacteria) within the genera *Rhodopila* ((s__Rhodopila sp903851415 and Rhodopila sp903915725, both MAGs recovered from ponds) and *Acidiphilum* (s__Acidiphilium sp028688905, recovered from a landfill in the North Eastern United States) encoded the entire *mmoXYZBD*. These members contained two copies of the *mmoX*, one clustering with hydrocarbon monooxygenases, while another *mmoX* sequence was similar to those of known methanotrophs (placed in the basal part of the "likely" *mmoX*, containing *Methylocystaceae*, *Methylococcaceae* and *Beijerinckiaceae*) (**Extended Data Figure 13**). Furthermore, all genomes encoded a *xoxF*-type methanol dehydrogenase, formaldehyde oxidation through the $H_4$MPT (Rhodopila genomes) or the GSH-linked pathway (*Acidiphilium* genome), formaldehyde assimilation through the $H_4$F, and an incomplete (*Rhodopila* 7/9, *Acidiphilium* 4/9) serine cycle. Additionally, carbon assimilation through the CBB cycle was encoded in all three genomes (**Extended Data Figure 1**). *Rhodopila globiformis*, a phototrophic purple bacteria isolated from a hot sulphur spring in Yellowstone, can utilise methanol and carbon monoxide[7], while *Rhodopila globiformis* and *Rhodopila sp. LVNP* have distinct lipid methylation signatures shared only by methanotrophic and acetic acid bacteria[8]. sMMO exhibits broad substrate specificity, oxidising aliphatic, aromatic, and halogenated hydrocarbons[9–11]. This might also apply to members of *Xanthobacteraceae, Rhodopila* and *Acidiphilium*, and assessment of methanotrophy would require cultivation or enrichment.

Three genomes within the family *Methylophilaceae*, order Burkholderiales, (GCA_013141125.1 s__JABFRO01 sp013141125 GCA_903926935.1 *Methylotenera* sp903926935, and GCA_947451235.1, *Methylotenera* sp947451235) encoded *mmoX*-like sequences, forming a distinct cluster grouping with other *Methylococcales mmoX* sequences. Furthermore, a *xoxF*-type methanol dehydrogenase, formaldehyde oxidation through the $H_4MTP$, and formaldehyde assimilation through the RuMP was encoded in all genomes (**Extended Data Figure 1**). Previous work have established synergistic relationships between methanotrophic *Methylobacter* and the denitrifying *Methylotenera* and shown genomic consensus for methylotrophy utilising xoxF and RuMP[12]. Another member of Burkholderiales (s__AVCC01 sp012927585, GCA_012927585.1) contained a *Methylotenera*-like sMMO operon. The key-enzymes *mxaFI* could not be identified, although the accessory genes *mxaACGJKL* were encoded. Furthermore, formaldehyde oxidation through the GSH-linked pathway and a complete CBB including RuBisCO was also present (**Extended Data Figure 1**).

*Methylophagaceae* members have been reported to lack methane monooxygenase, but encode PPQ methanol dehydrogenase, RuMP, formate dehydrogenase, and CBB cycle (contains RuBisCO)[13]. We were able to identify *pmoA* in s__GCA-002733105 sp014859375 (GCA_014859375.1), and *mmoX* in s__GCA-002733105 sp024636315 (GCA_024636315.1). Both genomes possess the genomic potential to oxidise methanol via *xoxF*-type methanol dehydrogenases, formaldehyde oxidation though the $H_4MPT$, while only the *pmoA*-containing genome encoded a complete RuMP pathway (**Extended Data Figure 1**). The *pmoCAB* gene were, however, placed on a short contig with no other genes, and could be a contaminating contig.

Within the genus *Candidatus* Macondimonas (Gammaproteobacteria) five genomes encoded putative *pmoA* (**Extended Data Figure 1**). *Ca.* M. diazotrophica have been shown to thrive in crude-oil contaminated coastal sediments, and genomic potential for methanotrophy along with hydrocarbon degradation and nitrogen fixation was proposed[14]. However, only a *pmoCAB* operon, along with methenyltetrahydrofolate cyclohydrolase (*fch* - $H_4F$ pathway), serine hydroxymethyltransferase (*glyA* - Serine cycle) and 5,10-methylenetetrahydrofolate reductase (MTHFR - folate metabolism)[14] were identified (**Extended Data Figure 1**). In one of the five genomes (sp021783685), *xoxF*, near-complete GSH-linked formaldehyde oxidation (*gfa, frmA*), and near complete CBB cycle (including RuBisCO genes but missing *glpx/fbp, rpiA/B*) could be identified. In the remaining four genomes, no genes encoding methanol nor formaldehyde oxidation could be identified. The serine cycle was mostly incomplete, and missing the key genes *sga* and *hprA*. The putative PmoA of *Ca*. M. diazotrophica was added to the GraftM package to improve classification of future potential and homologous sequences, but potential for methanotrophy within members of *Ca*. Macondimonas remains inconclusive.

Several species within the order Nevskiales were identified as potential methanotrophs, with *pmoA/amoA/emoA*-like sequences that grouped near to the particulate hydrocarbon monooxygenases (pHMOs) of *Cycloclasticus*[15] in a PmoA protein tree (**Figure 2a**). The genomic potential of *Solimonas aquatica*, CAU-1509 sp005047655 and *Panacagrimonas perspica* was investigated further, as these species had recently been proposed for methanotrophy[16]. Methanol oxidation (*xoxF*), formaldehyde oxidation via thiol dependent GSH-linked pathway (*gfa, frmAB*), and a nearly complete CBB pathway including both large- and small RuBisCO subunits (missing glpX/fbp) and several genes within the serine cycle (*mdh, glyA, ppc*) could be identified within *Solimonas aquatica* (RS_GCF_900111015.1) and *Panacagrimonas perspica* (RS_GCF_004364935.1) (**Extended Data Figure 1**). No methanol or formaldehyde dehydrogenases could be identified in CAU-1509 sp005047655 (RS_GCF_005047655.1). While the genomic potential of *Solimonas aquatica* and *Panacagrimonas perspica* indicate methanotrophy, CAU-1509 sp005047655 lacks essential pathways. Putative PmoA sequences from the three Nevskiales species were added to the GraftM package. In combination with the putative PmoA sequence from *Ca*. M. diazotrophica, they make up a distinct clade branching off next to the *Cycloclasticus* CuMMOs (**Figure 2a**). Classification within this cluster should be considered with caution.

Potential sMMO containing methanotrophs have also been suggested within the actinobacterial genus *Mycobacterium*[16]*.* Propane and ethane MOs share high sequence similarity with sMMO[17]. However, a recent study revealed the first experimental confirmation of methanotrophy within *Mycobacterium,* including downstream formaldehyde fixation by the RuMP pathway[18], and subsequently, another *Mycobacterium*, *Mycobacterium methanica MM-1*[19]*,* was isolated. The MmoX sequences of these newly described mycobacterial methanotrophs cluster with other mycobacterial non-sMMO sequences[19] (**Extended Data Figure 13**). Other mycobacterial genomes do harbour *mmoX*-like sequences similar to those non-sMMO monooxygenases but do not encode the potential to oxidise methanol or formaldehyde, and lack the methanotrophic carbon assimilation pathways (**Extended Data Figure 1**). Additionally, *mmoX* sequences were identified in a Chloroflexi MAG (GCA_012274105.1) from an Arctic cryosol methane SIP experiment[20]. However, this MAG was only 59% complete (checkM2), and is flagged as

”Undefined name (Failed Quality Check)” in GTDB, making interpretation of metabolic potential of little value.

Finally, we found *pmoA*-like sequences in the umbrella group, not falling within any well-described clades, within *Rugosibacter*, sp002842395, the UBA3067 (gammaproteobacteria), Proticoccaceae (either Nevskiales-type or the umbrella type), JADFDG01, DC-out160, f__Azotimanducaceae_A, g__JAJFNH01, g__JAAXHV01, g__UBA7887 and g__UBA6930. In all cases, metabolic potential for key methanotrophy modules were missing (**Extended Data Figure 1**). In all members of Burkholderiacea where we identified *pmoA*-like sequences, genes for methanol and formaldehyde oxidation were missing.

#### Supplementary Note 2: Elaborated annotation of recovered putative methanotrophic MAGs

To ascertain which of our recovered MAGs were putative methanotrophs, we created consensus metabolic models by investigating metabolic potential of all non-dereplicated MAGs. Genomes included in the analysis were filtered at >80% completeness and <10% contamination using CheckM1[21] and CheckM2[22], unless otherwise stated. The analysis was based on selected KOs assigned using DRAM, supplemented with additional HMMs. A list of KOs and their assignment to specific pathways can be found at **(Supplementary Data File 5, Supplementary Data File 6).** First, we evaluated the methane-centric metabolic potential by the presence of *pmo* and/or *mmo* operons, methanol oxidation through either *mxaFI*, and associated genes *mxaACGJKL* or *xoxF*. Next, we evaluated the oxidation of formaldehyde through either $H_4$MPT (*fae/fae-hps, mtdB, mch, ffsA, fhcA/B/C*) or GSH-linked (*gfa, frmA, frmB*) pathways. Then, we assessed the presence of carbon-assimilation modules for formate incorporation by $H_4$F (*fhs/ftfL*, *mdtA*, *fch*, *folD*) into the serine cycle (*glyA, sga, hprA, gck, eno, ppc, mdh, mtkA/B, mcl/aceB*). Alternatively, formaldehyde incorporation into the RuMP pathway (*hps, phi, pfb/pfk/fbp/glpX, fbaA/fbaB, tkt/tal*) was assessed. Finally, $CO_2$ fixation via the CBB cycle (*rbcL, rbcS, pgk, gapA, fbaA/fbaB, fbp/fbp3/glpX, tktAB/xfp/xpk, tktAB, rpiAB, prkB*) and oxidation of formate to $CO_2$ (*FDH*, *fdoGHD*) were analysed[23,24]. In addition to the methane-centric metabolism, we also investigated key genes involved in oxidation of hydrogen and carbon monoxide, nitrogen and sulphur conversions, storage of PHB, and types of terminal oxidases in the electron transport chain.

##### *Methylocapsa*

First, we explored the genomic potential of *Methylocapsa* members. Here, a clear split in the phylogenomic tree was observed between members encoding USCα *pmoA*, and members encoding either *mmoX* or *pmoA* sequences clustering outside of the USCα (more similar to those of *Methylocystis* and *Methylosinus* **Figure 2a**) (**Extended Data Figure 5**). *Methylocapsa* members encoded the MDH-*xoxF* for methanol oxidation, formaldehyde oxidation using the $H_4$MPT pathway (*ffsa, mch, fae, mtdb*), formate assimilation using the $H_4$F pathway (*fhs/ftfL* combined with either with *mdtA+fch* or *folD*[24]) and assimilation of carbon via the serine cycle (**Extended Data Figure 5**). Additionally, Methylocapsa MAG_7 and the associated species duplicates (except sp. dup. 10 and sp. dup. 22) encoded MDH-*mxaFI*. The CBB-cycle, including the large and small subunit of RuBisCO (*rbcL, rbcS*), was encoded in *Methylocapsa* MAGs 25, 26, 28, 35, 36, 41, 42, 43, and 44, along with the species duplicates of the same MAGs. The dephosphorylation of sedoheptulose 1,7-bisphosphate by SBAase (sedoheptulose-1,7-

bisphosphatase) (step 8/11) encoded by glpX-SEBP/fbp-SEBP (K11532, K01086, K01100) could not be identified (**Extended Data Figure 5**). However, FBA (Fructose 1,6-bisphosphate aldolase, K01624), encoded in *Methylocapsa spp.,* has been shown to be promiscuous working as both a FBPase and SBPase in the RuMP cycle of methylotroph *Bacillus methanolicus*[25]. Additionally, the regeneration part of the RuMP pathway can also be carried out by TA (transaldolase) variant instead of the SBPase variant[26]. Combined, this allows for alternative ways for putative $CO_2$-fixing *Methylocapsa spp.* to potentially carry out the CBB cycle. Furthermore, *Methylocapsa* members held the genomic potential for formate oxidation to $CO_2$ (**Extended Data Figure 5**).

In addition to the methane-centric metabolism encoded by *Methylocapsa* members, we also observed that the potential to oxidise carbon monoxide appeared to be conserved to a monophyletic clade of species and species duplicates within a cluster containing recovered MAGs for *Ca*. Lahnbergensis (MAG_16), *Methylocapsa* sp003162995 (MAG_25 and MAG_26), and MAG_17 and MAG_18 (**Extended Data Figure 6**). The potential to oxidise hydrogen using the Nickel iron (NiFe) group 1 hydrogenase appeared widespread, however incomplete (missing *hyaC*) in most members. The potential of these putative methanotrophic spp. to also oxidise hydrogen thus remains ambiguous. Few species (*Methylocapsa* MAG_35, MAG_41, MAG_43) encoded genes for NiFe group 2 and 3 hydrogenases. The genomic potential to fix nitrogen was encoded in a monophyletic clade consisting of *Methylocapsa* MAG_25, MAG_26, MAG_30, and *Methylocapsa* D3K7, and in most members of the clade with *Methylocapsa* MAGs 35-46.

All recovered members of *Methylocapsa* encode genes necessary for storage and utilisation of PHB (**Extended Data Figure 6**), as previously described in alphaproteobacterial methanotrophs[27]. Additionally, we wanted to investigate the complex IV of the electron transport chain, as the type of terminal oxidase encoded can be related to tolerance to sub-atmospheric levels of oxygen[28,29]. All MAGs, except the clade of *Methylocapsa* MAG_35 to MAG_46 encoded cytochrome c $aa_3$ (**Extended Data Figure 6**), classified as low affinity, thus requiring ambient oxygen-levels[28,29]. In contrast, the clade of *Methylocapsa* MAG_35 to MAG_46, along with *Methylocapsa* MAG_30 and its species duplicates, encoded the $cbb_3$-type cytochrome c oxidase with high-oxygen affinity, potentially allowing these *Methylocapsa* spp. to occupy habitats with sub-atmospheric levels of oxygen. Additionally, species found in wet habitats (*Methylocapsa* MAG_25, MAG_26, MAG_30) (**Figure 3a**) also encoded the high oxygen-affinity cytochrome bd (**Extended Data Figure 6**).

***Methylocystis***

Next, we investigated the metabolic potential of our MAGs recovered within *Methylocystis.* Here, members encoded either pMMO or sMMO, or both (**Extended Data Figure 5**). Differences in types of MMO encoded varied within members of the same species, such as the presence of *pmoCAB* and *mmoXYBZDC* in some species duplicates of MAG_27, which also encoded the *mxaF* type of the MDH in addition to MDH-*xoxF*. Additionally, *M.* silviterrae MAGs and species duplicates encoded the sMMO and the *mxaF* type of the MDH. The *mxaF* type of the MDH was also encoded in few non-sMMO harbouring *Methylocystis* MAGs, although this was sporadic. All members encoded formaldehyde oxidation using the $H_4$MPT pathway , and formate assimilation using the $H_4$F pathway (*fhs/fttL* and *mdtA*+*fch*), with detection of *folD* varying between species. Carbon assimilation was encoded via the serine cycle. No *Methylocystis* encoded a complete CBB-cycle or RuBisCO genes (**Extended Data Figure 5**).

Next, we evaluated additional metabolic potential in *Methylocystis* species. The ability to oxidise carbon monoxide was only found in *Methylocystis* MAG 31 and its species duplicate (**Extended Data Figure 6**). The potential to oxidise hydrogen was widely distributed amongst *Methylocystis* spp. with members encoding NiFe hydrogenase types 1, 2, and 3d. Presence of NiFe group 3d was restricted to one monophyletic clade of *Methylocystis* MAGs (MAG 27 to 43). The genomic potential to fix nitrogen gas was encoded in nearly all *Methylocystis* spp. Additionally, *hao* (hydroxylamine oxidoreductase), responsible for oxidation of hydroxylamine to nitrite, was encoded in the *Methylocystis* clade 1 (fen) abundant in the fens and sediments, and missing from clade 2 (bog) abundant in the bogs, molinia meadows and wet heaths, where we also identified the *pmoA2* (**Figure 4, Extended Data Figure 6**). *Methylocystis* members also held the potential to produce PHB, as previously observed in *Methylocystis rosea, M. parvus, M. SC2, M. hirsuta CSC1*[27]. The *scoA* and *scoB* (3-oxoacid-CoA transferase subunits A and B) are displayed as alternatives to the acetoacetyl-CoA synthetase (*acsA/aas*) in the degradation of PHB and conversion of acetoacetate and acetoacetyl-CoA[30], and are thus not missing for the synthesis of PHB in *Methylocystis.*

Most *Methylocystis* members encoded the high-affinity cytochrome bd in addition to the low-affinity $caa_3$-type cytochrome c oxidase (**Extended Data Figure 6**). Furthermore, two clades (MAG_27 to MAG_35 and associated species duplicates, along with MAG_18 and MAG 21) encoded a high-affinity cytochrome c $bb_3$ type. Overall, it appears that *Methylocystis* spp. held the genomic potential to utilise oxygen at sub-atmospheric concentrations.

***Rhodomicrobium***

We investigated the methane-associated metabolic potential of *Rhodomicrobium* species, where methanotrophy remains to be experimentally confirmed. We clearly observe that our recovered putative methanotrophic genomes cluster together in a phylogenomic tree, forming a monophyletic clade with members encoding putative pMMO and sMMO (**Extended Data Figure 10a**). Most *Rhodomicrobium* members, independent of the presence of putative MMOs, encoded the *xoxF*-type methanol dehydrogenase, GSH-linked formaldehyde oxidation, and CBB-cycle including RuBisCO (**Extended Data Figure 10a**). Additionally, the genomic potential for formate oxidation was encoded in most putative methanotrophic members. The putative methanotrophic *Rhodomicrobium* spp. encoded a high affinity $cbb_3$-type cytochrome c oxidase, potential to fix nitrogen (*nifHDK*), oxidise carbon monoxide (*coxSML*) and hydrogen ([NiFe] gr. 1, 2 and 3d), and potential to store PHB (**Extended Data Figure 10b**).

Putative methanotrophs within *Rhodomicrobium* also held the genomic potential to oxidise sulphur compounds, encoding *sat, aprAB, dsrAB, dsrC, dsrEFH* (**Extended Data Figure 10b**). As dissimilatory sulfite reductase can be both oxidative and reductive, we determined *Rhodomicrobium* DsrAB to be the oxidative type (reverse, rDsr) from protein phylogeny (**Extended Data Figure 9b**), using a DsrAB MSA downloaded from[2]. The transmembrane electron-transporting complex *dsrMKJOP* was encoded in only one of the MMO containing *Rhodomicrobium* (sp003153975, a MQ MAG from Stordalen Mire). Furthermore, the *dsrD* was absent, as expected from sulphur oxidisers[31]. The SOX complex was partially encoded (SOX ADXY).

**Gammaproteobacterial methanotrophs**

Our MAGs recovered within the gammaproteobacterial genera *Methyloglobulus*, UBA10906, *Methylovulum,* Methylobacter_C, JAGXGJ01, Methylobacter_A, *Methylobacter, Methylomonas* and CAIQWF01 encoded the expected general methane-centric metabolism (**Extended Data Figure 8a**). Methane oxidation was encoded by *pmoCAB* in most members, while *mmoXYBZDC* was encoded in UBA10906 MAG_6, Methylobacter_C MAG_14 and *Methylomonas* MAG_8, and *mmoXYBZC* in *Methylovulum* MAG_5. Methanol oxidation was encoded by MDF-*xoxF* in nearly all MAGs, while the presence of MDH-*mxaFI* varied. All MAGs encoded complete or near complete formaldehyde oxidation through the $H_4$MPT the pathway, and formaldehyde assimilation through the RuMP pathway. Additionally, most encoded the partial potential for formate oxidation to $CO_2$. The ability to oxidise carbon monoxide did not appear to be encoded in any of the gammaproteobacterial methanotrophic genomes (**Extended Data Figure 8b**). However, hydrogen oxidation was encoded in most genomes with the NiFe group 3d type hydrogenase (with the exception of only partially encoded in *Methyloglobulus* MAG_4 and missing in *Methyloglobulus* MAG_2). Additionally, NiFe hydrogenase group 1 and 2 was encoded sporadically across our recovered methanotrophic gammaproteobacterial MAGs. Fixation of $N_2$ was also encoded in most gammaproteobacterial methanotrophs, with presence varying between species within *Methyloglobulus*, UBA10906, and Methylobacter_C. The potential for partial denitrification to nitrous oxide (*narGHJI, nirK, norCB*) was encoded in *Methyloglobulus* MAG_3, Methylobacter_C sp002256465 MAG_17, Methylobacter_C MAG_11, 12, and 22 sp. dup. 1, JAGXGJ01, Methylobacter_A MAG 8 and 13, and *Methylobacter* MAG_6 and MAG_7, with *Methylobacter* MAG_6 encoding *narGHJI, nirS, norCB*, and *Methylobacter* MAG_7 encoding *narGHJI, norCB.* Assimilatory sulphate reduction was completely or partially encoded in most MAGs.

**USCγ**

We recovered MAGs from the genera USCg-Taylor and JACCXJ01 which encoded USCγ-like *pmoA*, and evaluated genomes with >80% completeness and <10% contamination (**Extended Data Figure 8a,b**), along with all (non-filtered) USCγ MAGs recovered in[32] and post oxidation rate experiments (**Extended Data Figure 8c**). All encoded a *pmoCAB* operon for methane oxidation (**Extended Data Figure 8a**). Methanol oxidation through MDH-*xoxF* was present in all our recovered MAGs except USCg-Taylor sp002007425 MAG_2, but could however be identified in a recovered genome from the same species (MFD05747.bin.2.8) (**Extended Data Figure 8c**). Formaldehyde oxidation through the $H_4$MPT pathway was conserved and complete in all MAGs but one (MFD06229.bin.1.238). Assimilation of formate by the $H_4$F pathway (*fhs/fttL* combined with either with *mdtA+fch* and/or *folD*) was encoded in most genomes, with completeness varying between MAGs. However, no complete pathway for carbon assimilation could be identified, as the key genes in the RuMP (hexulosephosphate synthase (*HxlA/hps)* K08093/K13812 and hexulosephosphate isomerase (*HxlB/phi)* K08094/K13812), serine cycle (glycerate dehydrogenase (*hprA)* K00018, glycerate-2-kinase (*gck)* K11529) and CBB (*rbcL, rbcS*) were not identified. Whether putative USCγ methanotrophs carry out carbon assimilation though the RuMP or serine cycles using alternative enzymes remains to be determined. Furthermore, USCγ MAGs encode *narGHJI*, with some members encoding *nirK* or *nirBD* (**Extended Data Figure 8b**).

Additionally, USCγ-related MAGs hold the genomic potential for PHB storage (**Extended Data Figure 8b**).
